# sabinaMBM: An R package for Multiscale Bayesian species distribution Modelling using INLA

**DOI:** 10.64898/2026.09.17.752384

**Authors:** Jennifer Morales-Barbero, Virgilio Gómez-Rubio, Javier Seoane, Antoine Adde, Teresa Goicolea, Rubén G. Mateo

## Abstract

1. Regional species distribution models (SDM) calibrated over spatially restricted extents tend to truncate species’ ecological niches. Existing nested SDM workflows integrate multi-scale information through sequential combination, which limits formal uncertainty propagation across scales and prevents regional predictions from being explicitly constrained within globally-informed niche boundaries.
2. We introduce sabinaMBM, an R package implementing joint multiscale Bayesian SDM within a unified probabilistic framework built on inlabru/R-INLA. It propagates uncertainty across scales without the computational bottlenecks of MCMC-based approaches. The framework offers multiple coupling architectures ranging from complete independence to hierarchical constraint that can be configured independently for intercepts and covariates.
3. In a range-margin population, hierarchical constraint most improves out-of-sample discrimination where regional data were scarcest, while leaving predictions unchanged where they already suffice, delivering gains precisely where sequential approaches are expected to struggle most. Applied to *Quercus petraea* across its Iberian trailing-edge, including a spatial field produced the largest single performance gain, consistent across every coupling configuration, and covariate responses diverged by scale for at least one climatic predictor. Under future climate, the constrained model yielded lower suitable habitat estimates and redistributed uncertainty in proportion to cross-scale agreement rather than uniformly.
4. sabinaMBM makes multiscale Bayesian inference accessible without specialist programming. This framework provides robust value for trailing-edge populations and spatial (invasive species) or temporal (climate change) projections where niche truncation risks ecologically implausible outcomes, while simultaneously delivering fine-resolution predictions with properly propagated uncertainty whenever global and regional covariates offer complementary information.

## 1 INTRODUCTION

Species distribution models (SDMs) are fundamental tools in biogeography, spatial ecology, and conservation, but their predictive accuracy depends on how they resolve the multi-scale nature of ecological drivers (Guisan & Thuiller, 2005; Keil et al., 2013; Mateo et al., 2017; Talluto et al., 2016). According to community assembly theory, species geographical distributions are shaped by hierarchical filters operating across a continuum of spatial scales (Lortie et al., 2004; Ricklefs, 2008), from broad-scale macroclimate constraints (hereafter “global”) to fine-resolution environmental factors such as microtopography and microclimate (hereafter “regional”) (Guisan & Rahbek, 2011). Spatially nested species distribution models (N-SDM) address this by combining regional-scale predictions within a globally-informed ecological context, overcoming the niche truncation and extrapolation bias of regional scale models (Chevalier et al., 2022; Goicolea et al., 2025; Guisan et al., 2025; Keil et al., 2013).

Recent tools such as N-SDM (Adde et al., 2026) and sabinaNSDM (Mateo et al., 2024) have operationalized this nested paradigm, combining global and regional correlative models through ensemble architectures (Araújo & New, 2007). These frameworks rely on sequential combinations, fitting each scale in a separate stage rather than jointly. They effectively quantify inter-algorithm and inter-fold disagreement when multiple algorithms or replicates are used, but this variability reflects differences in statistical methods, not across spatial scales. Nor are global and regional components estimated jointly, in part because fine-resolution covariates are seldom available across the broad extents (Moudrý et al., 2023) required to resolve global gradients and joint estimation is often computationally prohibitive/demanding at that scale (Tikhonov et al., 2020).

Joint multiscale Bayesian estimation offers an opportunity to overcome this limitation, estimating global and regional processes simultaneously and flexibly within a single likelihood so that they constrain one another directly, propagating uncertainty and information exchange across scales as a mathematical consequence (Mateo et al., 2019). Historically, Bayesian spatial inference faced severe computational bottlenecks with large datasets, the “big n” problem of Markov Chain Monte Carlo (MCMC) algorithms (Finley et al., 2007), —which had precluded fitting such joint architectures in practice. The stochastic partial differential equation (SPDE) approach (Lindgren et al., 2011) combined with integrated nested Laplace approximations (INLA; Rue et al., 2009) has since removed this barrier, rendering complex Bayesian spatial models computationally tractable. This tractability has been exploited to jointly integrate heterogeneous observation types (presence-only, presence-absence, counts) from multiple data sources —but still within a single spatial scale—, as in PointedSDMs (Mostert & O’Hara, 2023) and RISDM (Foster et al., 2024) Joint multiscale Bayesian SDMs have been demonstrated conceptually (Mateo et al., 2019) and calibrated for few species at landscape scale (Mateo et al., 2019), but to date, no dedicated software extends this joint estimation across spatial scales, explicitly coupling global and regional filters to address niche truncation within a unified probabilistic model.

To address this gap, we introduce sabinaMBM, an R package built upon the R-INLA and inlabru ecosystems (Bachl et al., 2019; Lindgren & Rue, 2015) to implement joint multiscale Bayesian SDMs. Conceptually, sabinaMBM shares the hierarchical motivation of nested SDM frameworks —integrating global and regional information— but formalizes this integration by jointly estimating global and regional processes within a single, flexible and probabilistic model. The package addresses multiscale modelling challenges through four core features: (1) native resolution of spatial misalignment across covariate resolutions (Alahmadi & Moraga, 2025; Barber et al., 2016; Gotway & Young, 2002); (2) support for binomial (presence-absence/background) and log-Gaussian Cox processes (presence-only) likelihoods; (3) flexible coupling architectures controlling cross-scale information flow; and (4) simultaneous global and regional spatial random fields to account for residual autocorrelation (Dovers et al., 2024; Gómez-Rubio et al., 2019; Palmí-Perales et al., 2021; Simpson et al., 2016). We detail the statistical architecture and coupling functions (Section 2), demonstrate the framework through a case study (Section 3), and discuss implications for multiscale ecological forecasting (Section 4).

## 2 THE sabinaMBM MODELLING FRAMEWORK

### 2.1 Joint multiscale Bayesian architecture

To integrate ecological processes across multiple spatial resolutions and extents, sabinaMBM implements a joint multiscale Bayesian framework (Box 1) using the R-INLA and inlabru packages (Bachl et al., 2019; Lindgren & Rue, 2008; Rue et al., 2009). Rather than fitting models in separate sequential stages as N-SDMs (Guisan et al., 2025), sabinaMBM simultaneously estimates broad-scale (“global”) and fine-scale (“regional”) processes (labels reflecting relative hierarchical position rather than absolute geographic extent). Coupling —the mechanism through which information estimated at the global scale is incorporated into, or used to inform, the regional process— governs this integration. Where a coupling architecture (see below) constrains the regional process against the global one, regional-scale predictions formally inherit uncertainty from the global scale as a direct mathematical consequence, rather than combining it after the fact.

### 2.2 Data preparation and variable selection

Essential pre-processing steps, including spatial thinning of species’ data and automated variable selection, are computed via the sabinaNSDM package (see, Mateo et al., 2024). Absence handling is addressed according to species’ data type. Presence-absence data supply true observed absences directly from the user’s occurrence dataset, modelled via a binomial likelihood. For presence-only data, sabinaNSDM’s formatting step automatically generates background points, but whether these are used depends on the likelihood family chosen for model fitting in sabinaMBM. Under the binomial approximation they serve as the available-environment component, whereas under the log-Gaussian Cox process (LGCP) approximation they are not used, and the intensity surface is instead integrated numerically over a discretization of the study area (the spatial mesh; Section 2.3) (for more details see, (Illian et al., 2012; Simpson et al., 2016).

### 2.3 Defining the spatial domain

The spatial domain of the analysis can be defined through a spatial mesh (i.e., a triangulation) that discretizes the study area, provides the spatial basis required to represent the continuous spatial random fields in the model, and provides the integration points used to evaluate the intensity surface under the LGCP likelihood. The construction of the spatial mesh via create_mesh() is mandatory for models incorporating spatial random fields (*S_s_*_ℎ*ared*_ and/or *S_RE_*, Section 2.4) or LGCP likelihoods, and not required otherwise (i.e., non-spatial binomial models). This function discretizes the domain into a Delaunay triangulation (Lindgren et al., 2011) from the processed nsdm.vinput object generated by sabinaNSDM. Users can refine the spatial mesh resolution, border extension, and domain masking through the edge, offset, and boundary.method arguments. Detailed parameterization guidelines and recommended default values are given in Supporting Information S1.2 and Table S1.3, respectively.

### 2.4 Model specification and inference

The multiscale Bayesian model comprises two coupled linear predictors: global (*η_GL_*) and regional (*η_RE_*):

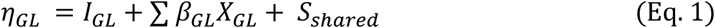

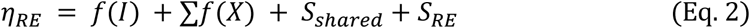

The global linear predictor follows a standard formulation. The global intercept (*I_GL_*) estimates the baseline occurrence intensity (log-Gaussian Cox process) or suitability (binomial) on the model scale, and the covariate coefficients (*β_GL_*) measure the average species’ response to global covariates. *I_GL_*and *β_GL_* are always estimated independently with weakly informative Gaussian priors, *N*(0, *σ*^2^).

The regional linear predictor (*η_RE_*) borrows strength from the global one through two coupling functions: *f*(*I*) for the intercept and *f*(*X*) for the covariate terms. Both act as mathematical bridges dictating whether the regional model estimates independent ecological effects, or inherits (conditions on) and locally refines the global knowledge (*I_GL_* and *β_GL_*). Both linear predictors additionally share a common broad-scale spatial field (*S_s_*_ℎ*ared*_) and, exclusive to the regional scale, a fine-scale field (*S_RE_*) capturing residual autocorrelation not explained by regional covariates, both detailed further below.

The model is fitted using the MBM.Modelling() function (Table 1), the main engine of sabinaMBM, which bridges the processed data to the MCMC-free inference machinery of R-INLA and inlabru (Bachl et al., 2019). Users specify family = ‘binomial’ for presence-absence/background data or family = ‘cp’ for a LGCP process fitted directly to presence-only records, with spatial integration resolved numerically over the spatial mesh instead of through background points (Illian et al., 2012; Lindgren et al., 2011). Supporting Information S1.3 details likelihood-specific interpretation and thinning considerations.

**Table 1.** Main functions and key arguments provided by the sabinaMBM package to configure and evaluate multiscale joint species distribution models.

| Function | Key arguments | Description |
| --- | --- | --- |
| `create_mesh()` | `edge`, `offset` | Discretizes the spatial domain using a Delaunay triangulation, balancing fine-scale resolution (`edge`) and boundary effect avoidance (`offset`). |
| `MBM.Modelling()` | `coupling.intercept` | Controls cross-scale flow of baseline suitability: NULL (regional-only), ‘unpooled’, ‘ordered_hierarchical’, or ‘bayesian_feedback’. Full specification in Supporting Information S1.2. |
|  | `coupling.covariates` | Defines how shared covariates are integrated across scales: NULL (regional-only), ‘unpooled’, ‘ordered_hierarchical’, ‘nested_shrinkage’, ‘scale_decomposed’, or ‘bayesian_feedback’. Full specification in Supporting Information S1.2. |
|  | `covariate.effects` | Allows flexible, per-variable specification of functional forms (‘drop’, ‘linear’ or ‘rw2’ splines with PC-prior complexity penalization). |
| | `*.pcprior.range/sigma` | PC priors governing the spatial ranges and variances of the broad-scale ($S_{shared}$) and fine-scale ($S_{RE}$) random fields. |
| `summary()` | `jmbm.inlabru` object | Returns structured model metadata, Bayesian fit criteria, hyperparameters, fixed/random effects, predictive performance, and cross-scale identifiability diagnostics. |
| `plot()` | `which` | Generates suitability and spatial-field prediction maps (‘pred’, ‘pred_Sre’, ‘pred_Sshared’) alongside diagnostic visualizations (e.g., ‘hyperparams’, ‘intercepts’, ‘correlogram’) from the `jmbm.inlabru` model object. |

### Coupling architectures

The coupling architectures define how information from the global model is shared with the regional model. MBM.Modelling function couples the intercept and covariate terms independently, offering three coupling architectures for the intercept *f*(*I*) and four for the covariates ∑*f*(*X*) (Supporting Information S1.4, Table S1.1), plus an option to remove cross-scale coupling entirely by setting the corresponding argument to NULL, available for both the intercept and the covariates.

For the intercept (coupling.intercept argument): 1) *f*(*I*) defaults to ‘unpooled’ (*I_RE_*, estimated independently of *I_GL_*). 2) ‘ordered_hierarchical’ shares the global estimate as the basis for the regional one (a mechanism we call a “hierarchical copy”), scaled by a freely estimated weight (*β_copy_*) whose prior, centred at one, expresses the expectation that regional and global baselines resemble each other unless the data indicate otherwise, without forcing an exact match. 3) ‘bayesian_feedback’ estimates it sequentially, using the global posterior as an informative prior for *I_RE_*, a scalable alternative for memory-intensive scenarios. Setting both coupling.intercept = NULL and coupling.covariates = NULL yields a regional-only single-scale model (useful for narrowly distributed or endemic taxa or as a null model); whereas setting NULL for only one component removes cross-scale coupling for that component while retaining it for the other.

For the terms shared between scales (coupling.covariates argument), ∑*f*(*X*) follows the same logic; scale-exclusive predictors are unaffected and enter each model independently. 1) ‘unpooled’ (default) estimates *β_RE_* independently of *β_GL_*; 2) ‘ordered_hierarchical’ instead applies the same hierarchical-copy mechanism used for the intercept (a per-variable *β_copy_*, freely estimated, scaling the global slope to produce the regional one); 3) ‘nested_shrinkage’ takes a different route, sharing the global slope exactly and adding a data-driven regional deviation (*δ_RE_*) shrunk toward zero, so the regional response departs from the global one only as far as the data support; 4) ‘scale_decomposed’ differs still further, partitioning each covariate into a global-scale trend and a regional-scale anomaly rather than constraining one against the other; and 5) ‘bayesian_feedback’ sequentially updates regional priors from global posterior moments. Setting coupling.covariates = NULL removes cross-scale coupling for scale-exclusive predictors. Regardless of whether spatial random fields are included, sabinaMBM evaluates covariates available at multiple, potentially mismatched resolutions (e.g., a coarse global layer and a fine regional layer for the same variable) directly at observation locations within a single joint likelihood, avoiding raster resampling except where the ‘scale_decomposed’ architecture explicitly decomposes a covariate across scales.

### Functional form of covariates

The covariate.effects argument defines the functional form of each covariate, globally or per variable: ‘linear’ by default, ‘rw2’ for second-order random walk smooth terms (Lindgren & Rue, 2008; Rue & Held, 2005) with smoothness governed by Penalized Complexity (PC) priors on the precision (Simpson et al., 2017; Supporting Information S1.5), or ‘drop’ to exclude a covariate at a given scale entirely.

Because coupling architectures constrain covariate coefficients directly, ‘rw2’ is only available for covariates that remain uncoupled between scales (Supporting Information S1.5). Non-linear responses other than rw2 splines (e.g., polynomial terms) are not additional covariate.effects options; instead, users can pre-compute them as derived covariates outside sabinaMBM and supply them as standard input covariates, which remain compatible with all coupling configurations.

### Spatial random fields

Missing covariates frequently induce residual spatial autocorrelation. To account for this residual spatial structure, sabinaMBM can include spatial random fields, which represent spatially structured variation in the response that is not explained by the used covariates (Václavík et al., 2012). When spatial fields are requested, continuous Gaussian random fields are additionally approximated as discrete Gaussian Markov Random Fields (GMRFs) over the spatial mesh built in Section 2.3, using the SPDE approach (Lindgren et al., 2011).

The package models spatial autocorrelation using two fields, which can be used individually or in combination: a broad-scale field shared across both scales (*S_s_*_ℎ*ared*_), and a fine-scale field (*S_RE_*) specific to the regional model (Dovers et al., 2024; Gómez-Rubio et al., 2019). *S_s_*_ℎ*ared*_ represents broad-scale spatial structure that may affect both global and regional predictors such as unmeasured biogeographical structure, major geographic barriers, and large-scale biotic gradients (e.g., (Boulangeat et al., 2012). *S_s_*_ℎ*ared*_ enters both global and regional predictors with a weight fixed at one (i.e., its contribution is not scaled by a separately estimated coefficient at either scale). Under’bayesian_feedback’, it is instead informed sequentially by the global posterior rather than re-estimated jointly, following the same logic applied above to the intercept and covariates (Supporting Information S1.6). Conversely, *S_RE_* captures fine-scale residual variance not explained by regional covariates, such as local topographic variation, fine-scale biotic interactions, spatial sampling bias, land use, or dispersal capacity) (Lenoir & Svenning, 2015; Renner et al., 2015). Unlike *S_s_*_ℎ*ared*_, it has no global counterpart.

Each field is activated by specifying its respective penalized complexity (PC) priors (arguments shared.pcprior.range, shared.pcprior.sigma, for *S_s_*_ℎ*ared*_ and regional.pcprior.range and regional.pcprior.sigma for *S_RE_*). The PC priors on the range specify the spatial range over which each field is expected to vary, while the priors on the sigma control the magnitude of the spatial variation. Omitting a field’s priors means it is not included in the model.

Including both spatial fields, allows distinguishing and interpreting separately ecological heterogeneity operating at different scales (Dovers et al., 2024; Mateo et al., 2017, 2019). However, they risk confounding each other’s spatial structure if their range parameters are numerically similar, causing double-smoothing and weak identifiability. To prevent this —what we refer to as adequate scale separation between *S_s_*_ℎ*ared*_ and *S_RE_*—, priors must be set so the two ranges remain distinct (Bakka et al., 2018; Simpson et al., 2017) (Supporting Information S1.6).

### Inference and validation

Inference of model parameters and their uncertainty rely on R-INLA deterministic approximations rather than MCMC sampling (for more details see Gómez-Rubio, 2020). The default inla.int.strategy = ‘eb’ uses Empirical Bayes estimation, that fixes hyperparameters at their posterior mode for speed, while ‘ccd’ integrates over the hyperparameter space when uncertainty quantification is critical. MBM.Modelling() evaluates model fit by combining predictive performance metrics (e.g., AUC, Brier score, log-score) with Bayesian criteria (e.g., WAIC, DIC). Model validation can additionally be performed via automated stratified k-fold cross-validation (cv.folds argument, default = 1, i.e. no cross-validation), disabled for LGCP since binary threshold-based metrics are not naturally suited to intensity-based point processes (Renner et al., 2015).

### 2.5 Model diagnostics and visualization

MBM.Modelling() actively monitors model behavior, issuing automated warnings for the most common diagnostic issues (among the full set detailed in Supporting Information S1.7) such as insufficient scale separation between the *S_s_*_ℎ*ared*_ and *S_RE_* fields (range ratio below); high posterior correlation between the two fields (above 0.7, indicating spatial redundancy; Dormann et al., 2013); and residual spatial autocorrelation (Moran’s I above 0.10, indicating unabsorbed structure). This built-in diagnostic system explicitly guides users through necessary structural refinements, such as adjusting PC priors or spatial mesh resolution.

A comprehensive summary() method delivers structured metadata, evaluation metrics, and parameter estimates, while plot() method generates diagnostic and visualization outputs via the which argument. Available outputs include posterior marginals, probability integral transform (PIT) histograms for calibration, isolated latent fields, and continuous prediction maps. The layer argument further maps predictive uncertainty [posterior standard deviation (‘sd’) or 95% credible interval bounds (‘q0.025’/’q0.975’) beyond expected suitability (‘mean’)], highlighting areas where data sparsity or cross-scale conflicts compromise model confidence.

## 3 ILLUSTRATIVE WORKED EXAMPLE

We apply the sabinaMBM framework to *Quercus petraea* (Matt.) Liebl. (sessile oak), a deciduous tree with a fragmented trailing-edge distribution at the southern margin of its Central European range. The Spanish dataset (N = 296 occurrences after spatial thinning) represents a fraction of the species’ global records (N = 1,115 after thinning) and occupies a climatically restricted subset of the full macroclimatic niche, missing 74% of the species’ global climatic conditions, though the regional distribution itself sits almost entirely within the global one (Fig. 1). Calibrating SDMs exclusively within this region constitutes a scenario where niche truncation is likely, making *Q. petraea* an appropriate case study for demonstrating hierarchical multiscale integration.

**Fig. 1.**
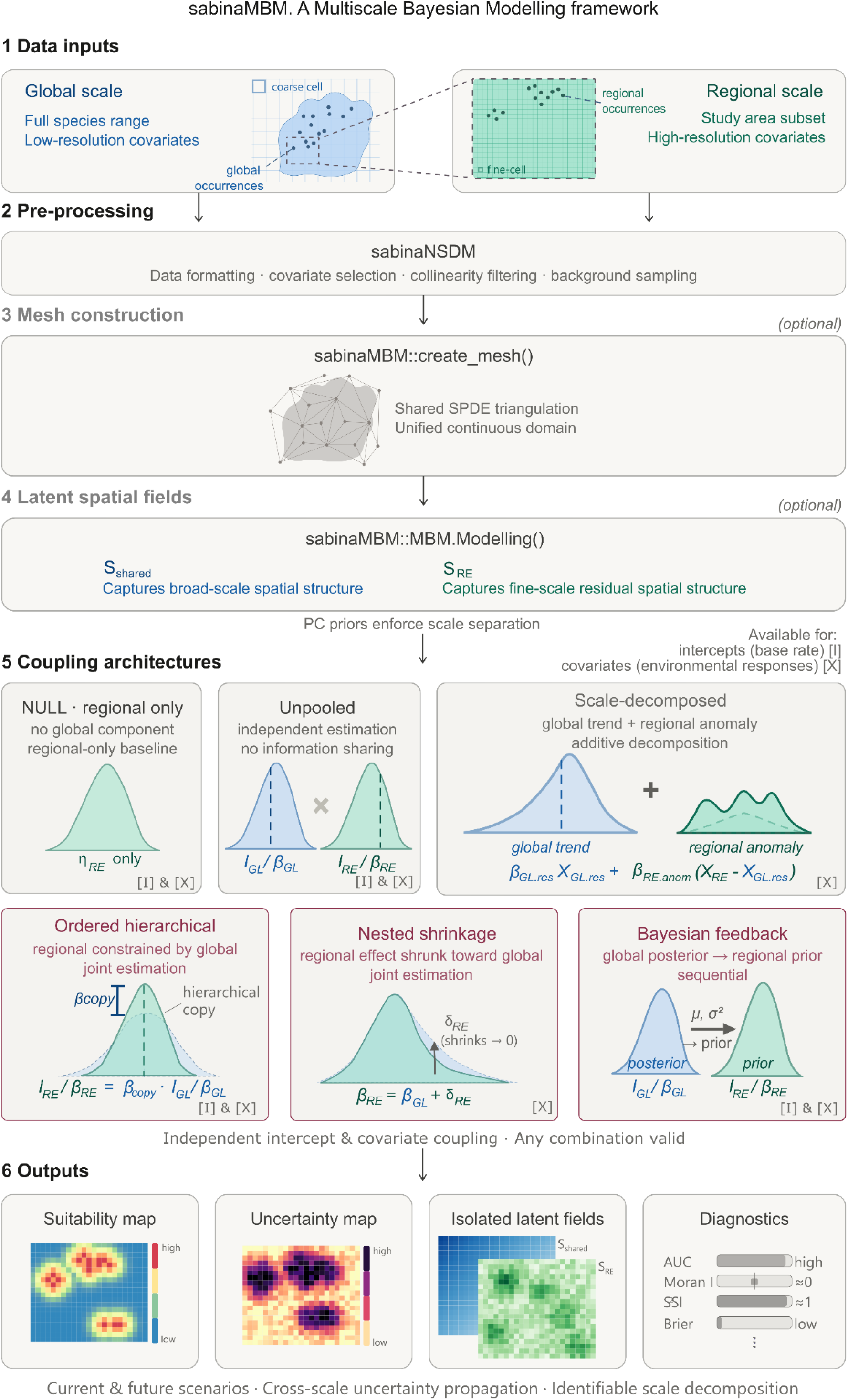
Niche overlap between global and regional environmental spaces of *Quercus petraea*. Points represent species occurrences projected onto the first two principal components of a PCA fitted on the combined dataset. Convex hulls delineate the occupied environmental space at each scale. Schoener’s D quantifies niche overlap; unfilling measures the proportion of the global niche absent from the regional space; stability measures the proportion of the regional niche contained within the global niche.

We used occurrence data and environmental covariates from the geoSABINA database (Goicolea et al., 2026), aggregated to global (∼50 km) and regional (∼10 km) resolution. Models were fitted under a binomial likelihood using 1,000 randomly sampled background points per scale. Alternative densities and sampling strategies were additionally evaluated as a robustness check (Supporting Information S4.1). All data pre-processing and covariate selection were executed through sabinaNSDM (Mateo et al., 2024). Using sabinaMBM, we fitted seven model configurations of increasing complexity (Table 2): a non-spatial regional-only baseline/null model, a spatial regional-only model with *S_RE_*, and five multiscale architectures (unpooled, ordered-hierarchical, scale-decomposed, nested-shrinkage, and Bayesian-feedback) integrating both *S_s_*_ℎ*ared*_ and *S_RE_*. All configurations were subsequently projected to a single GCM-SSP scenario for 2070 (MRI-ESM2.0, SSP585). Full reproducible code is in Supporting Information S3.

**Table 2.** Comparative predictive performance and computational cost across uniscale and multiscale model configurations. Coupling configurations are expressed as (coupling.intercept */* coupling.covariates). All spatial models include *S_RE_*; multiscale models additionally include *S_s_*_ℎ*ared*_. AUC, log-score, and Brier score are computed via the posterior predictive distribution on the training dataset and are comparable across all configurations; Moran’s I diagnose residual spatial autocorrelation. WAIC is reported for within-group comparison only, since likelihood dimensionality differs (N = 1,296 for uniscale and Bayesian-feedback configurations; N = 3,411 for the remaining multiscale configurations, including background points). Fitting time reports wall-clock duration on a standard laptop.

| Model (coupling<br>intercept/covariates) | N | WAIC | AUC | Log-score | Brier<br>score | Moran I | Fitting<br>time (min) |
| --- | --- | --- | --- | --- | --- | --- | --- |
| Non-spatial baseline<br>(NULL/NULL) | 1296 | 958.83 | 0.880 | -0.370 | 0.118 | 0.066 | 0.18 |
| Spatial regional-only<br>(NULL/NULL) | 1296 | 691.29 | 0.941 | -0.267 | 0.081 | -0.001 | 3.14 |
| Unpooled<br>(unpooled/unpooled) | 3411 | 1674.97 | 0.940 | -0.246 | 0.081 | -0.001 | 8.70 |
| Ordered-hierarchical<br>(ordered_hierarchical<br>/unpooled) | 3411 | 1656.98 | 0.940 | -0.243 | 0.081 | -0.003 | 24.83 |
| Scale-decomposed<br>(unpooled/scale_dec<br>omposed) | 3411 | 1645.10 | 0.947 | -0.241 | 0.077 | -0.001 | 7.67 |
| nested-shrinkage<br>(unpooled/<br>nested_shrinkage) | 3411 | 1667.76 | 0.941 | -0.245 | 0.081 | -0.001 | 10.22 |
| Bayesian-feedback<br>(bayesian_feedback/b<br>ayesian_feedback) | 1296 | 702.27 | 0.939 | -0.274 | 0.0821 | -0.001 | 17.86 |

Progressive model comparison (Table 2), evaluated in-sample, reveals a clear hierarchy of predictive improvement. The non-spatial baseline model achieved moderate discrimination (AUC = 0.880) with poor calibration (Brier = 0.118; log-score = −0.370). Incorporating *S_RE_* substantially improved both discrimination (AUC +7%) and calibration (Brier −31%, log-score −28%) over the non-spatial baseline —the largest single gain in the model comparison, consistent with residual spatial autocorrelation being the dominant source of misfit in the non-spatial baseline. Further integrating global-scale data through multiscale coupling yielded consistent additional log-score improvements over the spatial regional-only model (8-10.5% across joint configurations), with the exception of Bayesian-feedback, whose sequential architecture does not extend this log-score gain, though its discrimination and calibration remain comparable to the other configurations (Supporting Information S4.2).

We interpret the results below primarily through ordered-hierarchal, which most directly tests the ecological hypothesis motivating this case study —that in a trailing-edge, range-margin population, the regional niche is constrained by, rather than independent of, the global one, isolating this baseline-level mechanism from covariate-level coupling. Scale-decomposed scores marginally better on some metrics (Table 2), but tests a different hypothesis, partitioning each covariate into an additive global trend and regional anomaly instead. Supporting Information S4 details performance for the remaining model configurations. Out-of-sample, spatial cross-validation shows this constraint improves predictive performance over the regional-only model precisely where it is needed the most —evidence of transferability, not merely in-sample fit. Discrimination rises substantially in the data-poor trailing-edge region (Region 3: AUC 0.963 vs 0.693; Brier reduced by ∼70%), while the data-rich Region 2, where regional data are already sufficient, remains unaffected (Fig. 4; Supporting Information S4.5). This is the practical signature of hierarchical constraint. Where regional sampling is scarce, coupling lets the model borrow strength from the global fit, precisely the condition under which sabinaMBM’s joint architecture confers the greatest advantage.

Predictive performance is not the only signature of the joint multiscale Bayesian architecture. Spatial fields removed the strong residual autocorrelation present in the non-spatial baseline (Moran’s I: 0.066 → −0.001 to −0.003), with *S_s_*_ℎ*ared*_ and *S_RE_* achieving clear scale separation (posterior range 18.1° vs 2.5°) (Supporting Information S4.3). Environmental responses diverge by scale. Mean diurnal range (bio2) shows a strong, precise positive association globally (*β_GL_* = +0.267, 95% CI [0.112, 0.422]), but a regional estimate that shifts toward negative with a wide, largely uninformative credible interval (*β_RE_* = −0.067 [−0.486, 0.352]), hinting at a scale-specific response the global-only view would have missed (Fig. 2; Supporting Information S4.4). Under the future scenario, hierarchical constraint yields lower projected suitability than unconstrained regional model, both in overall magnitude (mean suitability 0.075 vs 0.089) and in extent (4.3% of Spain projected above the 0.5 suitability threshold versus 5.7% for the regional-only model). This reduction is consistent with the posterior coupling weight (*β_copy_* = 1.458, 95% CI [1.019, 1.895]), the estimated scaling factor linking the regional intercept to the global one (Fig. 3). Its credible interval excludes 1, indicating the regional baseline departs from a direct, unscaled copy of the global estimate. The future scenario reshapes uncertainty as well as suitability. Mean posterior standard deviation falls by 29.5% overall, and this reduction is not uniform across space but tends to concentrate in areas where the global prior most strongly revises suitability downward (cf. Figs 3C and 5C). Additionally, in regions where regional signal is sparse, the constraint stabilizes uncertainty instead of adding to it (Fig. 5; Supporting Information S4.8). This is direct empirical evidence of the formal cross-scale uncertainty propagation this architecture was designed to achieve.

**Fig. 2.**
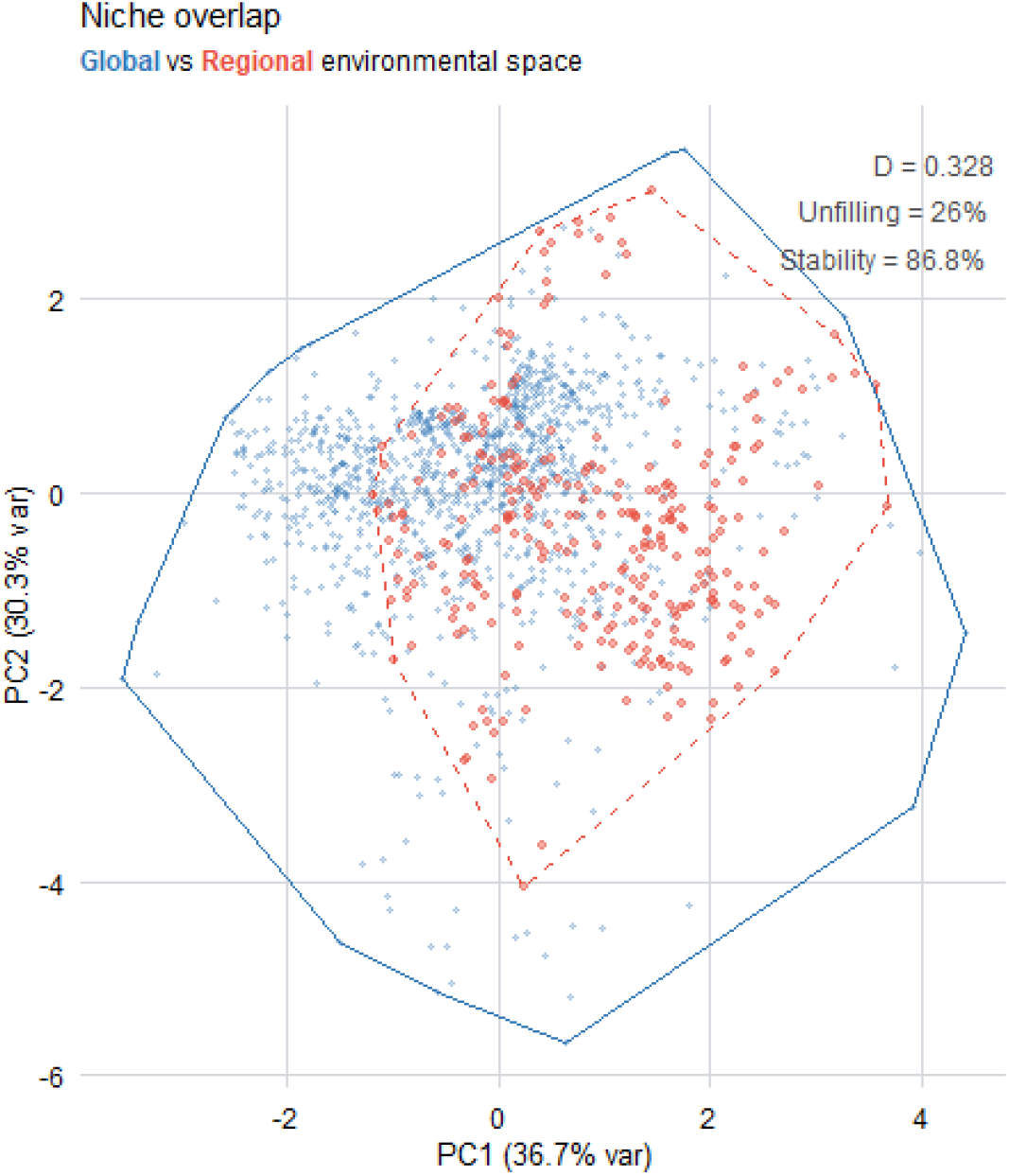
Scale-dependent covariate effects estimated by the ordered-hierarchical configuration (coupling.intercept = ‘ordered_hierarchical’, coupling.covariates = ‘unpooled’, with *S_s_*_ℎ*ared*_ and *S_RE_*). Global macroclimatic effects (*β_GL_*, blue circles) and regional effects estimated independently at each scale (*β_RE_*, red triangles) are shown with 95% credible intervals. Overlap of credible intervals with zero indicates covariates whose effect is not reliably distinguishable from zero at that scale.

**Fig. 3.**
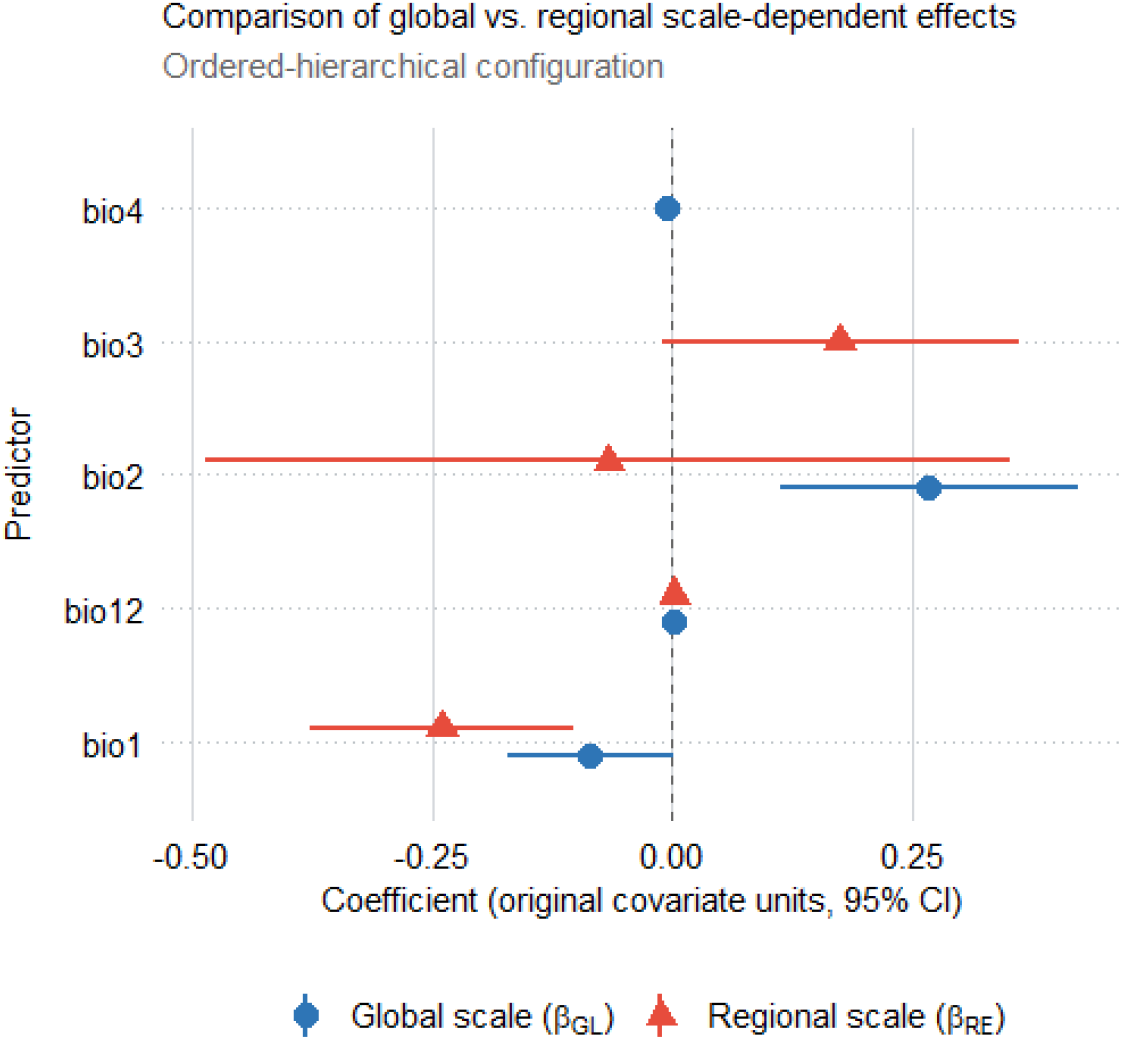
Cross-scale hierarchical coupling under future climate scenarios for *Quercus petraea*. Predicted habitat suitability from (A) the spatial regional-only (coupling.intercept and coupling.covariates = NULL, with *S_RE_*) and (B) the ordered-hierarchical model (coupling.intercept = ‘ordered_hierarchical’, coupling.covariates = ‘unpooled’, with *S_s_*_ℎ*ared*_ and *S_RE_*). (C) Pixel-wise difference (ΔSuitability = regional − hierarchical); warm colors indicate lower hierarchical suitability. (D) Prior (dashed) and posterior (solid) distributions of *β_copy_*; the dotted line marks the prior mean at 1.

**Fig. 4.**
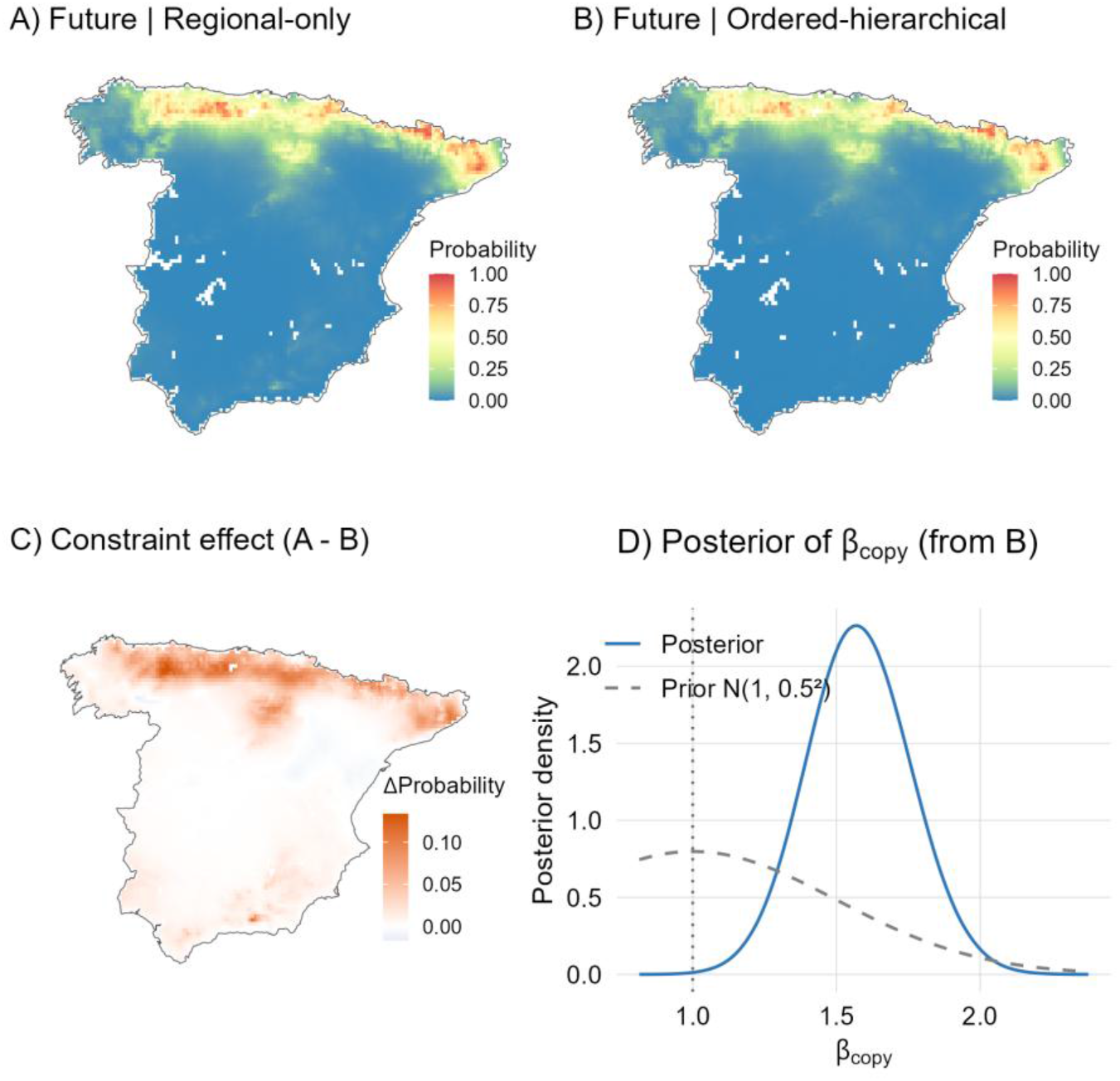
Spatial cross-validation by geographic region for *Quercus petraea*. Spain was partitioned into three non-overlapping regions defined by geographic coordinate thresholds. For each region, models were trained on the remaining two regions and evaluated on held-out records. The spatial regional-only (coupling.intercept and coupling.covariates = NULL, with *S_RE_*) and ordered-hierarchical configurations (coupling.intercept = ‘ordered_hierarchical’, coupling.covariates = ‘unpooled’, with *S_s_*_ℎ*ared*_ and *S_RE_*) are compared. ΔAUC is the difference in AUC (regional-only minus ordered-hierarchical). The Brier Skill Score (BSS) is the percentage improvement in Brier score of the ordered-hierarchical relative to the regional-only, 100 x (Brier_regional_-Brier_hierarchical_)/Brier_regional_; negative ΔAUC and positive BSS indicate that the hierarchical model outperforms the regional-only model. Dots represent occurrences used for model evaluation.

**Fig. 5.**
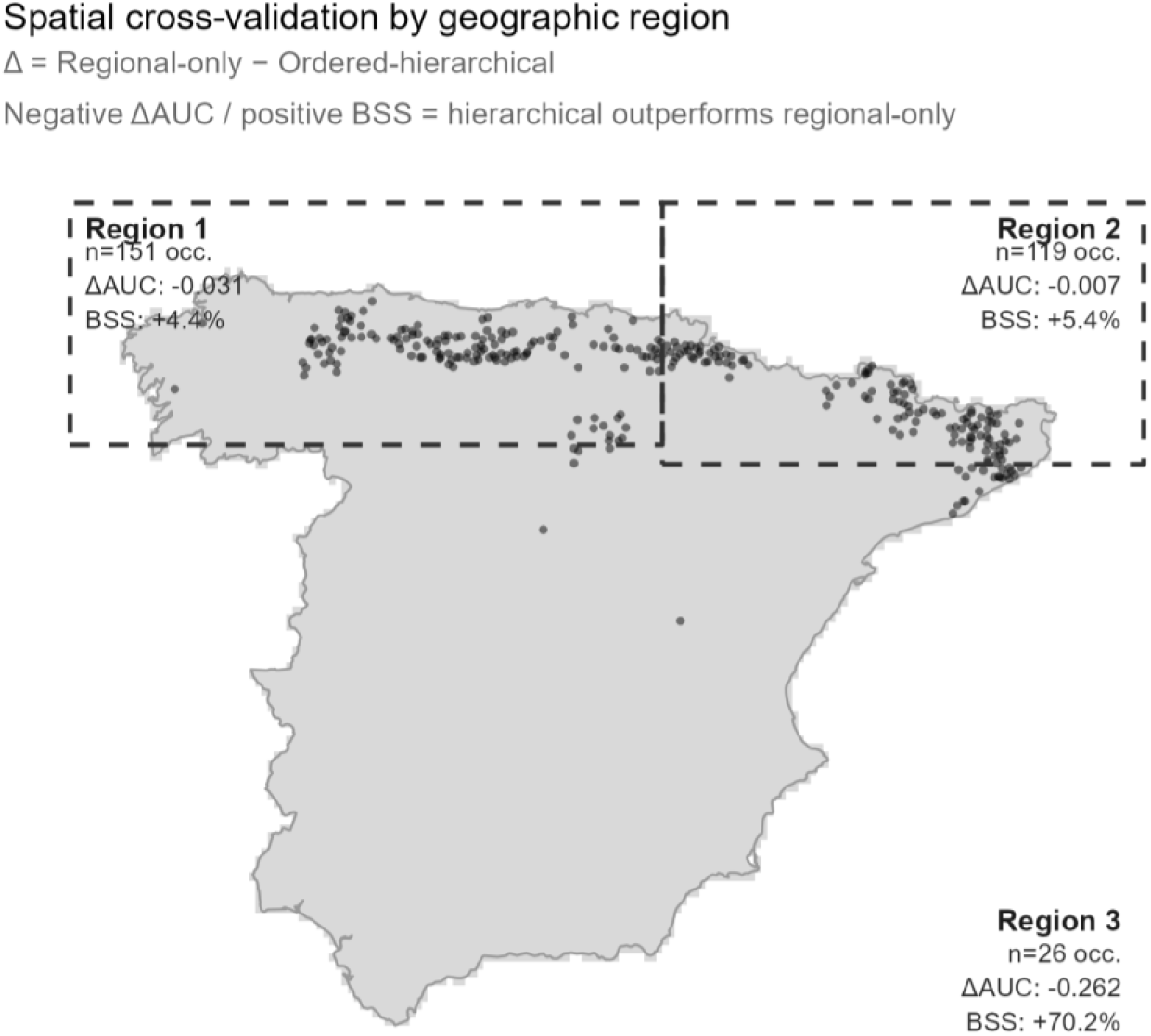
Predictive uncertainty and variance inflation under novel future climates for *Quercus petraea*. (A) Absolute predictive uncertainty (posterior standard deviation, SD) of the regional-only model, and (B) the ordered-hierarchical model (coupling.intercept =’ordered_hierarchical’, coupling.covariates =’unpooled’, with S_shared_ and S_RE_). (C) Pixel-wise (ΔSD = SD_regional_-SD_hierarchical_); warm colors indicate lower uncertainty under the hierarchical model, cool colors greater uncertainty.

## 4 DISCUSSION

sabinaMBM advances the nested SDM paradigm from a Bayesian perspective, formalizing cross-scale information exchange as a joint estimation problem in which hierarchical constraint and uncertainty propagation emerge directly from the shared likelihood. Leveraging the SPDE-INLA framework, the model integrates global and regional processes into a single sparse precision matrix, rendering multiscale Bayesian inference tractable without the MCMC bottlenecks (Bakka et al., 2018; Lindgren et al., 2011; Rue et al., 2009). In our case study, the most demanding coupling architecture (‘ordered_hierarchical’) fitted in under 25 minutes on a standard laptop (Intel Core i7-1165G7, 32 GB RAM), without specialized computing infrastructure. This architecture allows the regional model to genuinely learn from the global one, constraining regional predictions toward the globally-informed niche whenever regional data alone are not enough to determine them reliably, for example under niche truncation, sparse regional sampling, or extrapolation beyond the regional calibration domain. The posterior of *β_copy_*, the parameter that controls how closely the regional intercept copies the global one, illustrates this in practice, moving well away from a strict copy in our case study (mean = 1.578, 95% CI [1.225, 1.940]). Joint estimation also guarantees formal uncertainty propagation across scales in sabinaMBM’s fully joint architectures, since it keeps global and regional processes in one coherent posterior instead of passing credible intervals between separate model stages, where uncertainty risks collapsing (Figueira et al., 2024; Fletcher et al., 2019) —a risk with direct consequences whenever overconfident predictions in data-sparse regions could mislead conservation decisions.

A core strength of the package lies in its coupling architectures, which span a spectrum from independent to increasingly shared inference, allowing model structure to reflect alternative ecological hypotheses about how regional distributions and environmental responses relate to their global counterparts (Supporting Information S1.4 details the different options). ‘unpooled’ is appropriate when global and regional distributions or environmental responses may reflect distinct ecological contexts and should therefore be estimated independently, for instance when an insular population may differ substantially from its continental counterpart. Both ‘ordered_hierarchical’ and ‘nested_shrinkage’ allow the global response to inform regional estimation, which can be particularly valuable when niche truncation or scarcity of regional data is a genuine concern. Their underlying assumptions differ, however. ‘ordered_hierarchical’ encourages the regional response to follow the global pattern, while allowing its magnitude to vary, and is therefore appropriate when there are strong ecological grounds for expecting similar responses across scales; for example, when the regional study area encompasses only the range margin of a broadly distributed species. In contrast, ‘nested_shrinkage’ uses the global response as a reference while allowing the regional response to depart from it to the extent supported by the regional data, even reversing its direction; for example, when local ecological conditions may modify or override an environmental relationship observed across the species’ broader distribution. Our application illustrates the potential benefit of hierarchical information sharing. In the data-poor trailing-edge region, it substantially improved out-of-sample discrimination (AUC = 0.963 vs 0.693) and reduced the Brier score by ∼70%, while reducing uncertainty most strongly where regional evidence was weakest. Where regional sampling was already sufficient, the contribution of global information was comparatively small and the global and regional models converged, indicating that the practical gains from borrowing information across scales are concentrated where regional data are scarce. By informing these data-poor regions through the global niche, the framework replaces artificially wide intervals with estimates constrained by global ecological limits. The remaining architectures do not involve hierarchical coupling between global and regional responses. ‘scale_decomposed’ may be appropriate for ecological systems in which environmental variation operates differently across spatial scales, as it separates the response to broad-scale environmental variation from the response to local deviations (anomalies) around that broader environmental context. ‘bayesian_feedback’, in turn, provides a one-way transfer of information rather than joint coupling, using global estimates to inform regional inference in a subsequent modelling step. That is a potentially computationally efficient sequential alternative for large datasets. Finally, when information sharing across scales is unnecessary, the NULL option allows inference to be based entirely on regional data, as may be appropriate, for example, for endemic species.

Applying the framework carries specific assumptions and limitations. Estimating *S_s_*_ℎ*ared*_ and *S_RE_* together introduces identifiability risks addressed through explicit PC priors enforcing scale separation (Supporting Information S1.6), a decision that requires prior knowledge of the spatial grain of the processes being modelled. sabinaMBM evaluates covariates directly at observation coordinates (e.g., exact locations or atlas grid centroid). If records come as polygons instead of points (e.g., species checklists in grid cells), users must reduce them to a single coordinate, usually the centroid, since the package does not currently average covariate values across areas, a natural extension for a future release. ‘bayesian_feedback’ relies on a plug-in/sequential strategy for its two-stage architecture, which may underestimate posterior uncertainty in the second stage relative to full joint inference (Lee et al., 2025). It applies a first-order offset correcting for the presence-to-background ratio mismatch between global and regional datasets, though this correction is limited to the intercept and does not extend to covariate slopes. Users combining scales with markedly different sampling effort should still verify calibration explicitly. The package currently focuses on occurrence data, though its joint likelihood architecture provides a natural foundation for extensions to abundance data and multi-species joint distribution models.

As the availability of high-resolution biodiversity and environmental data continues to grow, the primary analytical challenge lies in rigorously integrating information across disparate spatial scales and extents, not only where niche truncation is a likely risk but wherever global and regional data offer complementary covariates (e.g., macroclimatic gradients combined with microclimate or microtopography). Beyond trailing-edge populations, this joint architecture is equally relevant for projections in space, such as invasive species forecasting into regions with no local calibration data (Guisan et al., 2025), and in time, such as climate change assessments where the regional model is by construction asked to predict conditions it never observed. In these cases, the value of sabinaMBM does not lie in outperforming single-scale models under current conditions, a gap our illustrative application shows can be modest, but in constraining and quantifying uncertainty precisely when and where extrapolation is unavoidable.

## 5 CONCLUSIONS

By automating SPDE-based spatial fields and hierarchical coupling within an MCMC-free framework, sabinaMBM bridges the gap between advanced Bayesian statistics and applied spatial ecology. The package advances the nested SDM paradigm by extending sequential combination toward a unified probabilistic flexible architecture for joint estimation, appropriate whenever a formal, mathematically guaranteed constraint between scales is required. Ecologically, this framework proves particularly valuable for trailing-edge populations and for forecasting under novel climate scenarios, contexts where the consequences of niche truncation are amplified (Chevalier et al., 2022; Goicolea et al., 2025; Guisan et al., 2025). Moving beyond mere methodological refinement, such unified architectures are essential to formally constrain regional predictions with global evidence, consistently propagate uncertainty, and deliver the biologically robust projections required for proactive ecosystem management.

## FIGURES AND TABLES

Box 1. The sabinaMBM modeling framework. A fully probabilistic R package for joint multiscale species distribution modeling using INLA, integrating global and regional occurrence data through a pipeline where users independently configure spatial mesh construction, latent spatial fields, and cross-scale coupling for both intercepts and covariates. Red boxes indicate coupling architectures that borrow strength from the global scale.

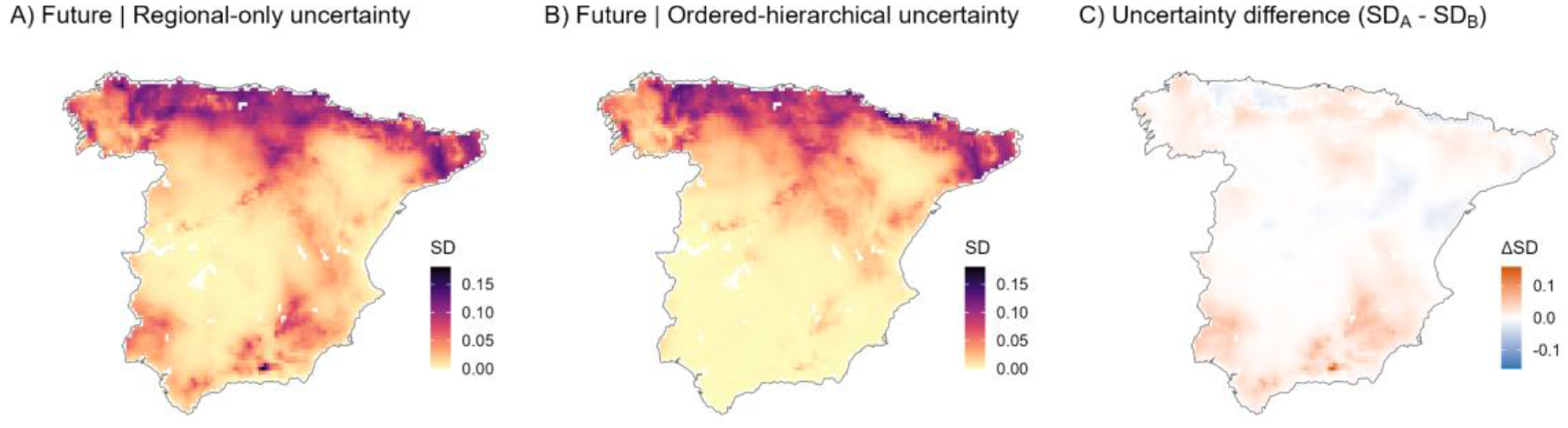

