## Supporting Information S1 for "sabinaMBM: An R package for Multiscale Bayesian species distribution Modelling using INLA"

**Supporting information S1: Extended methodology and mathematical formulation**

**Contents**

S1.1 Penalized complexity (PC) priors in multiscale models

S1.2 Defining the spatial domain and mesh resolution

Box S1.1 Example mesh construction workflow

S1.3 Likelihood-specific properties and adjustments

Box S1.2 Example thinning and likelihood specification workflow

S1.4 Mathematical specification of coupling architectures

Table S1.1 Coupling configurations available in sabinaMBM for the baseline intercept and predictors

S1.5 Non-linear covariate effects

Table S1.2 Compatibility of coupling.covariates architectures with covariate effect

S1.6 Prior specification for spatial random fields

S1.7 Post-fit diagnostics

S1.8 Quick-reference table for mesh and prior specification

Table S1.3 Quick-reference guide for user-configurable mesh and prior arguments

---------------------------

### S1.1 Penalized complexity (PC) priors in multiscale models

Hierarchical Bayesian architectures incorporating continuous spatial fields and non-linear covariate effects are highly flexible, but this flexibility risks overfitting and parameter unidentifiability. In models estimating multiple spatial scales simultaneously, traditional diffuse priors cannot effectively control variance partitioning among components. sabinaMBM therefore relies on PC priors (Simpson et al., 2017), which penalize deviations from a simpler base model unless data strongly support additional complexity, shrinking parameters toward parsimony via Kullback-Leibler divergence. Within sabinaMBM, PC priors govern two key components:

1. Non-linear covariate effects: The base model is a linear effect (zero spline variance). The PC prior on the rw2 precision penalizes unnecessary wiggliness, preventing overfitting to local environmental noise (Rue & Held, 2005).
2. Continuous spatial fields (SPDE): The base model is a spatially unstructured field (infinite range, zero variance). In multiscale architectures estimating $S_{shared}$ and $S_{RE}$ simultaneously, PC priors are strictly necessary to maintain identifiability and prevent double-smoothing (Bakka et al., 2018; Fuglstad et al., 2019).

#### S1.2 Defining the spatial domain and mesh resolution

The SPDE approach facilitates continuous spatial modelling, decoupling inference from arbitrary raster resolutions and extents. By partitioning the spatial domain into a Delaunay triangulation (mesh), this method transforms a dense spatial covariance matrix into a sparse Gaussian Markov Random Field (GMRF). This transformation reduces the computational complexity of spatial inference from $O(n^3)$ to $O(n^{3/2})$ (Lindgren et al., 2011), allowing sabinaMBM to perform simultaneous multi-resolution estimation without the computational bottlenecks of MCMC.

In practice, the mesh is defined (via `create_mesh()`) by a network of triangles with variable edge lengths and angles. Smaller triangles provide higher spatial resolution at greater computational cost. Users control this resolution via the `edge` argument (default c(0.5, 1), in CRS units), setting the maximum inner and outer triangle edge lengths. Methodological literature dictates that the maximum inner triangle edge length should be “several times smaller” than the expected spatial range to appropriately capture local structures without over-smoothing predictions. A general recommendation is to set this maximum triangle edge length to approximately 1/5 of the expected spatial range (Bakka et al., 2018; Dambly et al., 2023; Krainski et al., 2019) . Setting edge finer than the data can support, that is, generating substantially more mesh vertices than observations, risks a poorly identified, near-singular precision matrix in addition to unnecessary computational cost; users should verify that the number of mesh vertices remains proportionate to sample size, particularly for sparse regional datasets (Krainski et al., 2019).

Because the SPDE approximation assumes Neumann boundary conditions, spatial variance can be artificially inflated near mesh edges. The `offset` argument (default c(0.25, 0.5)), defines an outer buffer where this boundary artifact dissipates before affecting ecologically relevant predictions. Following formal recommendations, this outer extension must be set ≥ the expected spatial range to safely contain boundary effects (Anderson et al., 2025; Krainski et al., 2019).

The domain boundary itself is controlled via `boundary.method` argument, with three options: `‘convex_hull’` (default) and `‘concave_hull’` (requiring a concavity parameter controlling how tightly the hull wraps the points) derive the boundary directly from occurrence/background locations, while `‘raster_mask’` restricts mesh generation to environmentally valid space (e.g., non-NA predictor cells). The last option is strongly recommended for fragmented distributions or coastal/archipelago species, where it can substantially reduce computational burden by avoiding triangulation over open ocean or other non-habitat areas. Two further arguments apply exclusively to `‘raster_mask’`: `remove_holes` (default FALSE) removes internal habitat gaps before triangulation, and `proj.new.env` (default TRUE) automatically extends the mesh domain to cover future or novel environmental scenarios supplied by the user. Practical implementation is illustrated in Box S1.1.

**Box S1.1** Practical implementation of an optimal spatial mesh in sabinaMBM.

```
# Example: Creating a continuous mesh that protects against Neumann
# boundary variance inflation while strictly masking non-habitat areas
# (e.g., oceans).
# NOTE: edge and offset values must be given in the same spatial units as
# the coordinate reference system (CRS) of your environmental layers.

my_mesh <- sabinaMBM::create_mesh(
   nsdm_obj = my_sabinaNSDM_object,
   boundary.method = "raster_mask",
   edge = c(0.5, 1.5), # max triangle size: inner (0.5) and outer (1.5)
                       # domain,in raster units (here, degrees). Inner edge
                       # should be smaller than prior spatial range.
   offset = c(0.25, 0.5),  # outer buffer (>= expected spatial range) to
                           # absorb edge-effect variance inflation
   proj.new.env = TRUE,  # extend mesh for future climate scenarios
   plot = TRUE
```

```
)
```

#### S1.3 Likelihood-specific properties and adjustments

*Baseline intercepts in discrete approximations*

In discrete binomial approximations (`family = 'binomial'`) for presence-only data, the baseline intercept is intrinsically confounded with the unknown sampling effort and true species prevalence. Because presence-only data lacks true absences, estimating absolute occurrence probabilities is mathematically ill-posed without independent data on observation intensity (Fithian & Hastie, 2012). The unweighted binomial formulation in sabinaMBM is consequently designed to estimate relative habitat suitability (Renner et al., 2015). This relative ranking is mathematically robust to baseline prevalence shifts and correctly identifies environmental gradients and core habitats, but should not be interpreted as an absolute occurrence probability. When true presence–absence data are available instead, the observed 0s directly inform prevalence, so the fitted binomial probability is a valid, calibrated estimate of occurrence probability (Guisan & Zimmermann, 2000).

*Interpreting LGCP spatial intensity and thinning*

Discrete binomial models predict a bounded probability of occurrence (from 0 to 1) within an artificially defined spatial unit (e.g., a raster pixel). Continuous log-Gaussian Cox processes (`family = 'cp'`) instead operate in a continuous spatial domain, eliminating arbitrary grid discretization (Renner et al., 2015). The model output is the spatial intensity λ, defined as the expected number of presence records per unit of continuous area, strictly positive, without an upper bound of one (Aarts et al., 2012; Renner et al., 2015).

λ reflects the density of species reports, not necessarily the absolute abundance of biological individuals (Fithian & Hastie, 2012). Consequently, its ecological interpretation therefore depends heavily on the spatial thinning of occurrence records, controlled via `Min.Dist.Global` and `Min.Dist.Regional` arguments in `sabinaNSDM::NSDM.FormattingData()` (Mateo et al., 2024):

1. Thinned process (e.g., one point per pixels): λ estimates the intensity of occupied sites per unit area. This approach is recommended for opportunistic data (e.g., citizen science) to prevent the model from artificially inflating intensity due to oversampling. Functionally, it provides a robust index of relative habitat suitability.

2. Un-thinned process (negligible minimum distance): if researchers possess high-quality data where spatial aggregations are ecologically meaningful rather than artifactual, they can bypass thinning by setting a negligible minimum distance. In this scenario, λ evaluates the true spatial clustering of the occurrences, estimating the expected density of records per continuous unit area and preserving local abundance/aggregation patterns.

Regardless of the thinning strategy, the LGCP assumes occurrences are conditionally independent given λ(s). This assumption may be violated when spatial aggregation reflects true conspecific attraction or repeated detection of the same individual. Residual clustering should be assessed via `plot(x, which = "correlogram")` and `plot(x, which = "semivariogram")`. Practical implementation is illustrated in Box S1.2.

**Box S1.2** Implementation of spatial thinning in sabinaNSDM for LGCP models.

```
# Example A: Standard thinning (1 point per pixel) for biased data
# Output of 'cp' model = relative suitability (intensity of occupied sites)

myFormatting <- sabinaNSDM::NSDM.FormattingData(
   myInput,
   Min.Dist.Global = "resolution",
   Min.Dist.Regional = "resolution",
   ...
)

# Example B: NO thinning for high-quality data to preserve true clustering
# Output of 'cp' model = density of records (e.g., occurrences per km2)

myFormatting_dense <- sabinaNSDM::NSDM.FormattingData(
   myInput,
   Min.Dist.Global = 0.000001,
   Min.Dist.Regional = 0.000001,
   ...
)
```

**S1.4 Mathematical specification of coupling architectures**

*Baseline intercept (*`coupling.intercept`*)*

The intercept term, $f(I)$, governs the baseline suitability at regional spatial scale and controls how information flows from the global-scale to the regional-scale processes (Wikle & Berliner, 2005). In sabinaMBM, this flow is governed by the `coupling.intercept` argument, which offers four alternative coupling architectures of increasing hierarchical complexity (Table S1.1).

The default `'unpooled'` estimates the global and regional intercepts ($I_{GL}$ and $I_{RE}$) independently with weakly informative priors. This formulation imposes no hierarchical link between scales (i.e., the global intercept reflects broad-scale baseline suitability, while the regional intercept acts purely as an independent regional adjustment).

`'ordered_hierarchical'` implements an INLA adaptation of the multiscale ordered-hierarchical (MS-OH) framework (Zhou & Bradley, 2024). In this adaptation, the regional intercept is modelled as a true hierarchical copy of the global one via INLA's copy parameter. Following guidelines for weakly informative priors (Gelman et al., 2008) and INLA regularization (Simpson et al., 2017), the associated hierarchical scaling factor $\beta_{copy}$ is assigned a prior centred at unity ($N(1, 0.5^2)$). This acts as a Bayesian soft-constraint that reflects the prior expectation that regional baseline suitability resembles the globally-informed fundamental niche. Rather than treating the global prediction as a deterministic mask, this ordered-hierarchical prior imposes a stochastic constraint. This allows the regional posterior to deviate from global estimates when regional empirical evidence is sufficiently strong, implementing genuine cross-scale borrowing of strength (Chevalier et al., 2022; Zhou & Bradley, 2024). Crucially, constraining $\beta_{copy}$prevents the regional model from artificially inflating baseline suitability when calibrated on truncated, marginal regions.

`'bayesian_feedback'` is a sequential alternative. Instead of joint estimation, this configuration fits the global model first and uses its posterior distribution as an informative prior for the regional intercept, enforcing a unidirectional information flow. This updating-by-moments protocol (Figueira et al., 2024) provides a scalable alternative for computationally intensive scenarios (such as high-resolution grids or massive datasets) where simultaneous joint likelihood estimation might exceed memory or processing limits. While efficient, this method assumes an approximately Gaussian global posterior (Rue et al., 2009), a reasonable assumption for well-identified intercepts, but a potential limitation if the global posterior is strongly skewed. The transferred intercept mean is further adjusted by a first-order offset correcting for the presence-to-background prevalence ratio mismatch between the global and regional datasets. This correction is limited to the intercept and does not extend to covariate slopes.

`coupling.intercept = NULL` fits a regional-only model that excludes the global component entirely. This configuration can serve as a null model for assessing the global component's contribution, or for applications where only regional-scale inference is required or available (e.g., narrowly distributed or endemic taxa) (Guisan et al., 2025).

*Predictor coefficients (*`coupling.covariates`*)*

The term $\sum f(X)$ defines how covariate effects are shared, modified, or estimated independently across scales. To address spatial misalignment challenges (Barber et al., 2016), sabinaMBM introduces the `coupling.covariates` argument. For any covariate shared across scales, this argument offers five architectures (Table S1.1):

`'unpooled'` (default) estimates regional coefficients ($\beta_{RE}$) independently of global $\beta_{GL}$. This allows the regional model to be fully data-driven by local observations without sharing information across scales, which is appropriate when species are expected to respond differently to global and regional environmental gradients.

Conceptually inspired by multiscale spatial decomposition (Cressie & Wikle, 2015), the `'scale_decomposed'` architecture partitions each shared covariate into a global-scale trend and a regional-scale anomaly, both estimated on a common standardization scale, so that $\beta_{GL.res}$ $and$ $\beta_{RE.anom}$ are directly comparable. This formulation is ecologically motivated by the premise that species may respond differently to broad-scale geographical gradients versus fine-scale regional anomalies (Suggitt et al., 2011). This is the sole configuration that internally resamples the global raster to the regional resolution, since computing the anomaly requires subtracting the two at matching pixels; the global raster must therefore be of equal or coarser resolution than the regional one.

`'ordered_hierarchical'` implements a true hierarchical copy (Zhou & Bradley, 2024), structurally identical to the intercept mechanism, applied to a covariate coefficient rather than a baseline. By integrating global-scale uncertainty directly into $\beta_{copy}$'s posterior, this genuine cross-scale borrowing of strength constrains regional responses to remain consistent with globally estimated environmental relationships, reducing the risk of ecological over-extrapolation into non-analogous regional climates, while allowing data-driven departures when regional evidence supports divergence.

`‘nested_shrinkage’` implements an additive hierarchical shrinkage by partitioning the ecological response rather than the environmental data. It parameterizes the regional effect as the shared global coefficient plus a data-driven regional deviation. This deviation is governed by a PC prior that shrinks its variance toward zero, effectively pulling the regional effect toward the global trend unless local data provide clear evidence of divergence. Unlike the multiplicative constraint in `‘ordered_hierarchical’`, this configuration learns the strength of cross-scale borrowing entirely from the data, without centering on a fixed a priori target.

Moving away from the joint-likelihood framework used elsewhere, `‘bayesian_feedback’` implements a sequential, unidirectional information flow. After harmonizing shared covariates into a unified global Z-space, the global model is fitted first and injects its posterior moments ($\mu_{GL}$, $\sigma^2_{GL}$) as the Gaussian prior for $\beta_{RE}$. sabinaMBM adapts this updating-by-moments protocol from Figueira et al. (2024) —originally developed to combine different sampling processes at a single spatial resolution to the multiscale context. This sequential approach is particularly advocated because joint likelihood estimation can be numerically unstable for scales with large information disparities. The approach is pragmatic but restrictive, transferring information only through the first two moments, leaving higher-order posterior features unpropagated, assumes approximate Gaussianity of the global posterior (Rue et al., 2009; flagged automatically when |skewness| > 1), and is structurally limited to linear covariate effects. `'bayesian_feedback'` must be applied to the intercept and every shared predictor simultaneously; it cannot be combined with any other coupling architecture within the same model call.

`coupling.covariates = NULL` removes any cross-scale coupling, allowing scale-exclusive predictors. All intercept and predictor configurations are summarized in Table S1.1.

**Table S1.1** Coupling configurations available in sabinaMBM for the baseline intercept and predictors. The joint linear predictor is $\eta_{GL}$ and $\eta_{RE}$ (Eq. 1 and 2), with $f(I)$ and $\sum f(X)$ coupling functions detailed below. $I_{GL}$ and $I_{RE}$ are the global and regional intercepts; $\beta_{GL}$ and $\beta_{RE}$ the covariate coefficients; $\beta_{copy}$ the hierarchical copy weight; $\tilde{X}_{GL.res}$ is the global covariate resampled to regional resolution, and $(X_{RE} - \tilde{X}_{GL.res})$ its regional anomaly, with responses $\beta_{GL.res}$ and $\beta_{RE..anom}$ respectively. $\hat{\mu}_{GL}$ *and* $\hat{\sigma}^2_{GL}$ are the global posterior mean and variance used in sequential updating. $I_{GL}$ and $\beta_{GL}$ are always estimated independently with weakly informative Gaussian priors $N(0,\sigma^2)$. * Mean further adjusted by a first-order prevalence-offset.

| Configuration (arg) | Mathematical form \| Prior specification | Information flow | Recommended use | Main caveat |
|---|---|---|---|---|
| **A. BASELINE INTERCEPT (`coupling.intercept`)** | | | | |
| `NULL` | $f(I) = I_{RE}$ ; Prior: $I_{RE} \sim N(0,\sigma^2)$ | None — regional only | Regional-only inference | No cross-scale information |
| ‘`unpooled`’ (default) | $f(I) = I_{RE}$ ; Prior: $I_{RE} \sim N(0,\sigma^2)$ | Independent, no hierarchy | Distinct global/regional baselines | No borrowing across scales |
| ‘`ordered_hierarchical`’ | $f(I) = \beta_{copy} I_{GL}$ ; Prior: $\beta_{copy} \sim N(1, 0.5^2)$ | Joint hierarchical copy | Range-margin/subset populations (peripheral or disjunct regional populations) | Single $\beta_{copy}$; weak if regional data scarce |
| ‘`bayesian_feedback`’ | $f(I) = I_{RE}$ ; Prior: $I_{RE} \sim N(\hat{\mu}_{GL}, \hat{\sigma}^2_{GL})$ * | Sequential (GL → RE) | Large datasets | One-way; no full uncertainty propagation |

**B. COVARIATES (`coupling.covariates`)**

| | | | | |
|---|---|---|---|---|
| `NULL` | $f(X) = \beta_{RE}X_{RE}$ ; Prior: $\beta_{RE} \sim N(0, \sigma^2)$ | None — global excluded | Regional-only covariate effects | Excludes NULL-set covariate(s) from the global submodel |
| `‘unpooled’ (default)` | $f(X) = \beta_{RE}X_{RE}$ ; Prior: $\beta_{RE} \sim N(0, \sigma^2)$ | Independent, no hierarchy | Substantially different responses | No shared information |
| `‘scale_decomposed’` | $f(X) = \beta_{GL.res}\tilde{X}_{GL.res} + \beta_{RE.anom}(X_{RE} - \tilde{X}_{GL.res})$; Prior: $\beta_{GL.res}, \beta_{RE.anom} \sim N(0, \sigma^2)$ | Additive trend/anomaly split | Disentangling broad trend from local anomaly | Depends on resampling accuracy |
| `‘ordered_hierarchical’` | $f(X) = \beta_{copy}\beta_{GL}X_{RE}$ ; Prior: $\beta_{copy} \sim N(1, 0.5^2)$ | Joint hierarchical copy | Range margins, multiplicative effect | Per-predictor version of the intercept risk |
| `‘nested_shrinkage’` | $f(X) = (\beta_{GL} + \delta_{RE})X_{RE}$ ; Prior: $\delta_{RE} \sim N(0, {\sigma_\delta}^2)$ with PC-prior on $\sigma_\delta$ | Joint, data-learned shrinkage | Strength learned from data | Extra shrinkage hyperparameter to tune |
| `‘bayesian_feedback’` | $f(X) = \beta_{RE}X_{RE}$ ; Prior: $\beta_{RE} \sim N(\hat{\mu}_{GL}, \hat{\sigma}^2_{GL})$ | Sequential (GL → RE) | Large datasets | Same one-way flow, per predictor |

**S1.5 Non-linear covariate effects**

In sabinaMBM, species' responses to environmental gradients can be modelled flexibly using second-order random walk splines (rw2; (Lindgren & Rue, 2008; Rue & Held, 2005) (Table S1.2). However, highly parameterized splines can easily overfit the data by tracking localized, spurious noise in the environmental predictors rather than true physiological limits. To prevent this, spline smoothness is governed by PC priors on the precision.

Within the covariate.effects argument, users control this flexibility via two hyperparameters, u and alpha, such that $P(\sigma > u) = \alpha$ (Simpson et al., 2017). By default, `‘rw2’` effects `u=0.5`, `alpha=0.01`, expressing only a 1% prior probability that the curve's structural variation exceeds 5 standard deviations of the (Z-standardized) covariate. This default specification expresses a strong prior belief that the species' response is smooth. The spline will only deviate into more complex shapes if the data provide overwhelming evidence for such non-linearities, keeping the estimated ecological niche biologically plausible.

`‘rw2’` integrates seamlessly with `‘unpooled’` and `‘scale_decomposed’`, but is not supported with `‘nested_shrinkage’`, `‘ordered_hierarchical’`, and `‘bayesian_feedback’` (Table S1.2). For `‘bayesian_feedback’`, a multi-parameter spline's posterior cannot be meaningfully summarized or transferred as a single Gaussian prior mode. For `‘nested_shrinkage’` and `‘ordered_hierarchical’`, although INLA's copy construct can technically reference a non-linear component, scaling an entire spline curve by a single hierarchical weight would assume the regional response shares the exact shape of the global curve, differing only in amplitude (an assumption at odds with the ecological rationale for allowing non-linear responses in the first place). sabinaMBM therefore restricts these three mechanisms to linear global effects by design, not due to a limitation of copy itself.

**Table S1.2** Compatibility of coupling.predictors architectures with covariate functional form. coupling.intercept is unaffected, since the intercept has no functional-form choice.

| **coupling.covariates** | **linear** | **rw2 (spline)** | **Reason** |
|---|---|---|---|
| `NULL` | ✓ | ✓ | No cross-scale coupling to constrain |
| `‘unpooled’` | ✓ | ✓ | Independent estimation, no constraint |
| `‘scale_decomposed‘` | ✓ | ✓ | Trend/anomaly split integrates seamlessly |

| | | | |
|---|---|---|---|
| `‘ordered_hierarchical‘` | ✓ | X | Copy scales an entire curve; assumes matching global/regional shape |
| `‘nested_shrinkage‘` | ✓ | X | Same constraint as ordered_hierarchical |
| `‘bayesian_feedback‘` | ✓ | X | Multi-parameter posterior cannot be summarised as a single Gaussian prior |

**S1.6 Prior specification for spatial random fields**

Jointly estimating a broad-scale shared field ($S_{shared}$) and a fiend-scale regional field ($S_{RE}$) introduces the risk of weak identifiability. If both operate at similar spatial resolutions, they confound each other, producing double-smoothing artifacts. sabinaMBM addresses this through PC priors on both spatial range (ρ) marginal standard deviation (sigma σ) for each field (Fuglstad et al., 2019), shrinking each toward a structureless base model (infinite range, zero variance) unless data provide compelling evidence otherwise. Both fields are NULL by default; only $S_{RE}$ may be specified alone, $S_{shared}$ without $S_{RE}$ is not permitted.

To choose biologically meaningful ρ and σ bounds, a standard heuristic SPDE assumes an expected ecological range of ~20% of the spatial domain's maximum diameter (Barber et al., 2016). Following the simulation-based optimal calibration demonstrated by Fuglstad et al. 2019), we then set the prior range lower limit (ρ) of $S_{RE}$ to 1/10 of this expected spatial range. To guarantee identifiability and scale separation, $S_{shared}$'s expected range must be significantly larger (at least 3× that of $S_{RE}$); when the sigma ratio ($\sigma_{shared}/\sigma_{RE}$) exceeds 1.5 (Blangiardo & Cameletti, 2015); or when posterior field correlation exceeds 0.7. When any of these thresholds is exceeded, users should revise their PC prior specification before interpreting model outputs.

A common intuition is to structure $S_{RE}$ as a hierarchical refinement of $S_{shared}$ ($S_{RE} = S_{shared,resampled} + \delta_{RE}$). This is not recommended within the INLA/inlabru framework for three reasons. First, SPDE fields are GMRFs whose precision matrices encode conditional independence between spatial nodes (Lindgren et al., 2011); a deterministic dependency between the two fields breaks this Markovian structure and can bias posterior marginals (Bakka et al., 2018). Second, resampling $S_{shared}$ from a coarse spatial mesh onto a finer regional mesh introduces bilinear interpolation error that contaminates the fine-scale signal the decomposition aims to isolate (Barber et al., 2016). Third, R-INLA does not natively support hierarchical copy operations over continuous SPDE fields, making such a structure technically impractical without substantial custom development (Rue et al., 2009).

The additive, conditionally independent decomposition adopted in sabinaMBM, where $S_{shared}$ and $S_{RE}$ enter as separate additive components estimated jointly from their respective likelihoods, resolves these issues while preserving genuine multiscale inference. Cross-scale information flows through $S_{shared}$ entering both the global and regional likelihoods simultaneously with a fixed unit weight (Dovers et al., 2024; Gómez-Rubio et al., 2019) (‘`bayesian_feedback`’'s sequential architecture is the exception, detailed below) capturing shared biogeographic autocorrelation at a broad scale. $S_{RE}$ captures residual fine-scale heterogeneity unique to the regional scale. Scale separation is achieved statistically through the priors above, not through a deterministic hierarchy (Fuglstad et al., 2019; Simpson et al., 2017). Under `'bayesian_feedback'`, $S_{shared}$ follows the same logic as the intercept and covariates. The global posterior informs the regional prior. $S_{shared}$ is fitted once from the global likelihood as a SPDE field with PC prior. Its posterior mean range and sigma are then used to build a new, median-matched PC prior for the field's regional-likelihood fit.

#### S1.7 Post-fit diagnostics

Even with carefully specified PC priors, it is crucial to verify that the joint multiscale model successfully resolved the spatial fields and environmental responses without confounding parameters. sabinaMBM automatically computes diagnostics table (`mod$Summary$Diagnostics`), printed via `summary(mod)`, flagging seven conditions worth checking routinely:

1. Scale separation: range ratio ($S_{shared}/S_{RE}$); a ratio closer to 1 means both fields are absorbing the same variance (Bakka et al., 2018).
2. Variance balance: sigma ratio ($\sigma_{shared}/\sigma_{RE}$) should stay below 1.5; higher values suggest the shared field is outcompeting regional-scale structure (Blangiardo & Cameletti, 2015).
3. Field independence: posterior correlation between $S_{shared}$ and $S_{RE}$ should stay below 0.7; higher values indicate overlapping structure.
4. Residual autocorrelation: Moran's I of regional residuals should stay below 0.10; higher values mean the spatial field(s) failed to absorb true autocorrelation.
5. Informative data: a fixed effect's posterior should differ meaningfully from its diffuse prior; near-identical values mean the data said little about that covariate.

6. Hyperparameter identifiability: a hyperparameter's credible-interval-to-median ratio should stay below 25.
7. Predictive reliability: fewer than 1% of observations should show a failed or unreliable conditional predictive ordinate (CPO).

Beyond these automated checks, users should also visually inspect posterior non-linear response curves and spatial fields, since overly wiggly environmental responses or highly fragmented spatial fields point to insufficiently tight PC priors (lower `u`/`alpha` in `covariate.effects`). Any flagged condition should prompt revisiting the corresponding prior specification, mesh resolution, or coupling architecture before interpreting model outputs.

#### S1.8 Quick-reference table for mesh and prior specification

**Table S1.3** Quick-reference guide for specifying mesh and prior arguments in sabinaMBM. Values assume a projected coordinate reference system measured in metres (e.g., UTM).

| **Component** | **Argument** | **Recommended value (rationale)** |
|---|---|---|
| **in `create_mesh()`** | | |
| Mesh resolution (inner) | `edge` | "several times smaller" (e.g., ≤1/5) of expected regional patch size (Dambly et al., 2023). |
| Mesh boundary (outer) | `offset` | ≥ expected regional range (patch size), to absorb boundary variance inflation (Anderson et al., 2025). |
| **in `MBM.Modelling()`** | | |
| Broad-scale range | `shared.pcprior.range` | ≥3× `regional.pcprior.range` (Bakka et al., 2018). |
| Fine-scale range | `regional.pcprior.range` | ~1/10 of expected regional patch size (Fuglstad et al., 2019). |
| Regional variance | `regional.pcprior.sigma` | Bound σ to limit spatial probability shifts (Fuglstad et al., 2019). |
| Global variance | `shared.pcprior.sigma` | Smaller than `regional.pcprior.sigma`, heavily penalized, so $S_{shared}$ acts as a soft biogeographic |

| | | |
|---|---|---|
| | | background without double-smoothing regional structure (Fuglstad et al., 2019). |
| Non-linear covariates | `covariate.effects` | Default `u=0.5`, `alpha=0.01`. P(σ > 0.5 log-odds) = 1% (Simpson et al., 2017). |
