## Supporting Information S2 for "sabinaMBM: An R package for Multiscale Bayesian species distribution Modelling using INLA"

#### Supporting Information S2: A quick-start guide

7 References 15

8 Session information 15

---

### 1 Overview

This quick-start guide demonstrates the core workflow of **sabinaMBM**: fitting joint multiscale Bayesian species distribution models that integrate broad-scale (global) and fine-scale (regional) occurrence data with spatial random effects via INLA.

We model the distribution of *Quercus petraea* across two scales (Europe and Iberian Peninsula), first with a **non-spatial baseline** and then with the full **joint** multiscale model (coupled global and regional scales with hierarchical coupling). Additional examples show how to fit non-linear covariate effects and log-Gaussian Cox process formulations.

For complete documentation, see the main article (Section 2) and `?MBM.Modelling`.

### 2 Installation

```
# INLA from dedicated repository
install.packages("INLA",
  repos = c(getOption("repos"), INLA = "https://inla.r-inla-download.org/R/stable"),
  dep = TRUE)

# sabinaNSDM (data preparation)
remotes::install_github("geoSABINA/sabinaNSDM")

# sabinaMBM
remotes::install_github("JMoralBa/sabinaMBM")
```

```
library(terra)
library(patchwork)
library(inlabru)
library(INLA)
```

---

### 3 Data Preparation

#### 3.1 Load species occurrences and environmental rasters

```
# Species identifier
SpeciesName <- "Quercus.petraea"

# Occurrence records at two spatial scales
data(Quercus.petraea.xy.global, package = "sabinaMBM")   # European extent
data(Quercus.petraea.xy.regional, package = "sabinaMBM") # Iberian Peninsula

# Environmental rasters (current)
data(expl.var.global, package = "sabinaMBM")
data(expl.var.regional, package = "sabinaMBM")
expl.var.global   <- terra::unwrap(expl.var.global)
expl.var.regional <- terra::unwrap(expl.var.regional)

# Future scenario
data(new.env, package = "sabinaMBM")
new.env <- terra::unwrap(new.env)
```

#### 3.2 sabinaNSDM pipeline: formatting and covariate selection

The sabinaNSDM package (Mateo et al., 2024) handles background sampling, spatial thinning, and covariate selection before model fitting. The function NSDM.InputData() accepts optional pre-built background data frames via Background.Global and Background.Regional if a custom sampling strategy is preferred. To run a regional-only model, set spp.data.global = NULL and expl.var.global = NULL.

```
## INPUT DATA PREPARATION
myInput <- sabinaNSDM::NSDM.InputData(
  SpeciesName       = SpeciesName,
  spp.data.global   = Quercus.petraea.xy.global,
  spp.data.regional = Quercus.petraea.xy.regional,
  expl.var.global   = expl.var.global,
  expl.var.regional = expl.var.regional,
  new.env           = list(new.env),
  new.env.names     = "Scenario1"
)

## DATA FORMATTING & THINNING & BACKGROUND GENERATION
myFormatting <- sabinaNSDM::NSDM.FormattingData(
  myInput,
  nPoints           = 1000,
  Min.Dist.Global   = "resolution",
  Min.Dist.Regional = "resolution",
  save.output       = FALSE
```

```r
)

## SELECT COVARIATES
mySelvars <- sabinaNSDM::NSDM.SelectCovariates(
  myFormatting,
  corcut            = 0.7,
  algorithms        = c("glm"),
  save.output       = FALSE
)
```

---

#### 4 Non-spatial baseline (optional)

This section fits a classical, regional-only model, purely as a point of comparison. It's optional. Skip straight to Section 5 if you want to fit the joint multiscale model directly.

A classical SDM fitted exclusively to the regional dataset. It evaluates covariates but ignores spatial autocorrelation and broad-scale niche limits entirely.

```r
mod_baseline <- MBM.Modelling(
  jmbm_obj              = mySelvars,
  family                = binomial(link = "logit"),
  spde.mesh             = NULL,        # No spatial fields
  coupling.intercept    = NULL,        # No hierarchical architecture
  coupling.covariates = NULL,
  proj.new.env          = FALSE
)
```

##### 4.1 Suitability map baseline model

```r
plot(mod_baseline, which = "pred", layer = "mean")
```

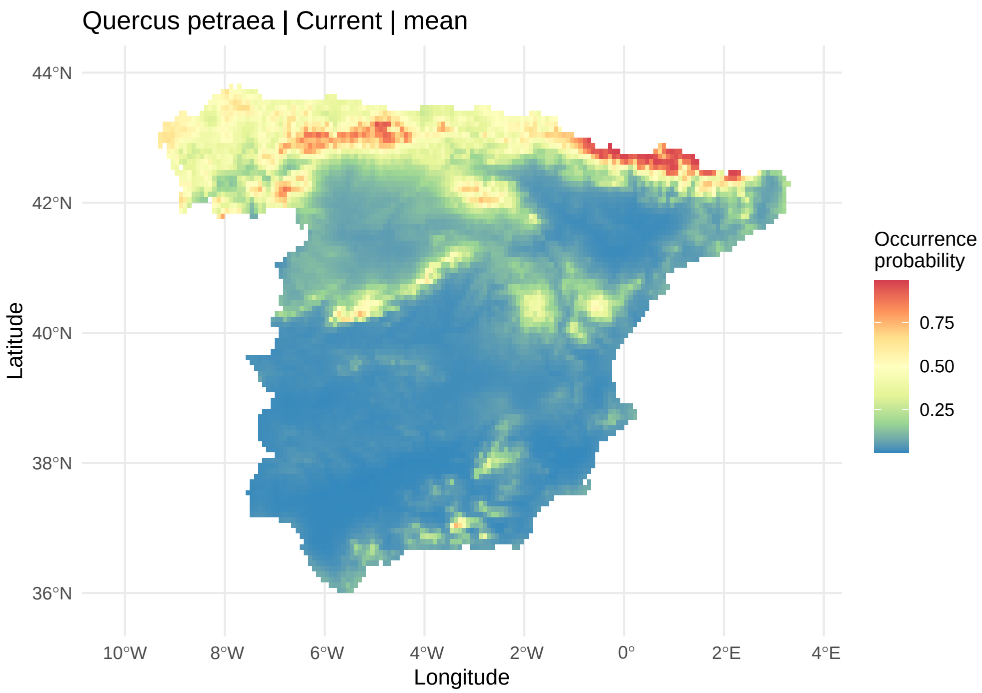

**Fig. S2.1.** Current suitability (posterior mean) under the non-spatial baseline model.

This is the kind of model you'd fit without sabinaMBM's multiscale machinery: no spatial field, no information from the global scale. sabinaMBM's own approach (Section 5) adds both at once.

---

#### 5 The sabinaMBM approach: joint hierarchical model

This is sabinaMBM's core workflow: a joint multiscale model that couples global and regional scales through spatial random fields and a hierarchical intercept/coefficient constraint. Building it takes three steps: a spatial mesh, PC priors for the spatial hyperparameters, and the model fit itself.

##### 5.1 Create spatial mesh

The mesh breaks the spatial domain into a network of triangles, which is how INLA approximates spatial autocorrelation efficiently instead of working with a dense covariance matrix. `edge` controls triangle size (inner/outer domain) and `offset` controls how far the mesh extends beyond the data boundary, both in the units of the raster CRS (degrees for geographic coordinates, km for projected coordinates). A finer inner mesh (smaller `edge[1]`) improves spatial resolution at the cost of increased computation.

```
myMesh <- create_mesh(
  nsdm_obj          = mySelvars,
  edge              = c(2, 10),
  offset            = c(1, 5),
```

```
  boundary.method = "raster_mask",
  plot            = TRUE
)
```

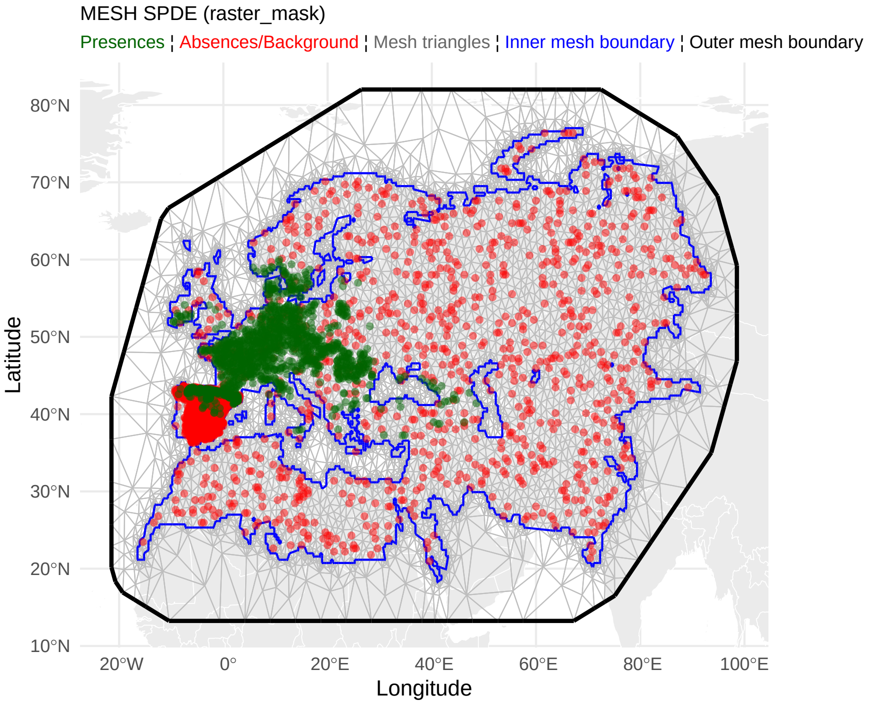

#### 5.2 Define PC priors for spatial hyperparameters

PC priors penalise unnecessarily complex spatial structure unless the data provide strong evidence for it. This is what keeps $S_{shared}$ and $S_{RE}$ statistically identifiable from one another.

```
# Practical range (distance where correlation   0.1)
# Tail probability: P(range < u) = alpha
regional.pcprior.range <- c(2, 0.01)      # Regional: ~2° range
regional.pcprior.sigma <- c(1, 0.01)      # Spatial SD

shared.pcprior.range   <- c(5, 0.01)      # Broad-scale: ~5° range
shared.pcprior.sigma   <- c(1, 0.01)
```

#### 5.3 Fit the joint hierarchical model

Relative to the non-spatial baseline (Section 4), the joint model adds two things together: spatial random fields (a broad-scale $S_{shared}$ and a fine-scale $S_{RE}$, both defined using the mesh and priors above), and a hierarchical constraint where the regional model borrows strength from the global baseline suitability via INLA's hierarchical copy mechanism.

```r
mod_hierarchical <- MBM.Modelling(
  jmbm_obj               = mySelvars,
  family                 = binomial(link = "logit"),
  spde.mesh              = myMesh,
  regional.pcprior.range = regional.pcprior.range,
  regional.pcprior.sigma = regional.pcprior.sigma,
  shared.pcprior.range   = shared.pcprior.range,
  shared.pcprior.sigma   = shared.pcprior.sigma,
  coupling.intercept     = "ordered_hierarchical",
  coupling.covariates    = "ordered_hierarchical",
  proj.new.env           = TRUE
)
```

The summary reports model fit, hyperparameters, random and fixed effects, and predictive and diagnostic performance

```r
summary(mod_hierarchical)
```

```
##
## ======================================================================
##  Summary for: Quercus.petraea
## ======================================================================
##
## ------ Model metadata ------
##                  Field                          Value
##          Species name:                Quercus petraea
##            Model type: MBM: Sre + Sshared + covariates
##         Family | Link:                binomial | logit
##   Coupling (Intercept):           ordered_hierarchical
##  Coupling (Covariates):                       unpooled
##
## ------ Model fit (Bayesian criteria) ------
##                              Metric    Value
##                                 DIC 1674.500
##                                WAIC 1656.980
##    Marginal log-likelihood (log ML) -972.240
##                 LCPO (sum log-CPO) -828.870
##  MLPD (mean log predictive density)   -0.243
##
## ------ Hyperparameters ------
##                                   Parameter        Value
##      Sre field: Range (posterior mean ± SD)  2.49 ± 1.15
##      Sre field: Sigma (posterior mean ± SD)  1.74 ± 0.41
##  Sshared field: Range (posterior mean ± SD) 18.06 ± 2.96
##  Sshared field: Sigma (posterior mean ± SD)  2.05 ± 0.21
##          Precision for IGlobal (mean ± SD)  0.38 ± 0.17
##
```

```
## ------ Random effects ------
##                                                Term
##                         IGlobal (mean ± SD, CI95%)
##  IRegional (copy from IGlobal, mean ± SD, CI95%)
##        Copy   (IGlobal -> IRegional) (mean ± SD)
##                                      Value
##  -4.73903 ± 0.5475 (-5.81211, -3.66594)
##  -7.43376 ± 0.85882 (-9.11702, -5.7505)
##                                1.58 ± 0.18
##
##   Note: IGlobal / IRegional: scale-specific intercepts (log-odds scale).
##  Copy   (IGlobal → IRegional): scaling of the hierarchical intercept copy (ordered_hierarchi
##
## ------ Fixed effects ------
##   [Global scale]
##     coef  estimate       sd      2.5%     97.5% signif       tail_p
##   bio1GL -0.085622 0.044167 -0.172187  0.000944        0.0522255798
##  bio12GL  0.000990 0.000382  0.000241  0.001739    *** 0.0095759705
##   bio2GL  0.267290 0.079085  0.112286  0.422294    *** 0.0007141565
##   bio4GL -0.005003 0.001434 -0.007813 -0.002193    *** 0.0004603175
##
##   [Regional scale]
##     coef  estimate       sd      2.5%     97.5% signif      tail_p
##   bio1RE -0.240666 0.069867 -0.377602 -0.103730    *** 0.000549058
##  bio12RE  0.001689 0.000584  0.000544  0.002833    *** 0.003838267
##   bio2RE -0.066949 0.213824 -0.486036  0.352139        0.752584816
##   bio3RE  0.174565 0.094840 -0.011317  0.360448        0.065901995
##
##   Note: GL = global scale; RE = regional scale.
##   Coefficients are back-transformed to original covariate units (effect per unit of X).
##
## ------ Predictive performance ------
##                                 Metric     Value
##                       AUC (full model) 0.9401182
##  Tjur R² (discrimination coefficient) 0.5190000
##                            Brier score 0.0810000
##                      Brier Skill Score 0.5400000
##                                   RMSE 0.2850000
##    Observed-predicted correlation (r) 0.7360000
##
## ------ Diagnostics ------
##                                                Metric  Value
##                                    Residual Moran's I -0.003
##                 Max CI/median ratio (hyperparameters)  1.961
##   Scale-separation ratio (range_Sshared / range_Sre)  7.266
##           Variance ratio (sigma_Sshared / sigma_Sre)  1.177
##                     Sshared-Sre field correlation (r)  0.067
##                         Variance explained by Sre (%) 42.994
```

```
##                    Scale separation Index (SSI) [0-1]  0.850
##  Posterior field redundancy r²(S_RE, S_shared) [0-1]  0.004
```

Look at the Copy β (covariateGL → covariateRE_oh) rows in the Random effects block above. That suffix marks the hierarchical copy constraint linking each regional coefficient to its global counterpart. A value close to 1 means the regional response is echoing the global one; a value far from 1, or with a wide credible interval crossing 0, means the regional data are pulling away from the global pattern for that covariate.

##### *5.3.1 Suitability map hierarchical model*

```
plot(mod_hierarchical, which = "pred", layer = "mean")
```

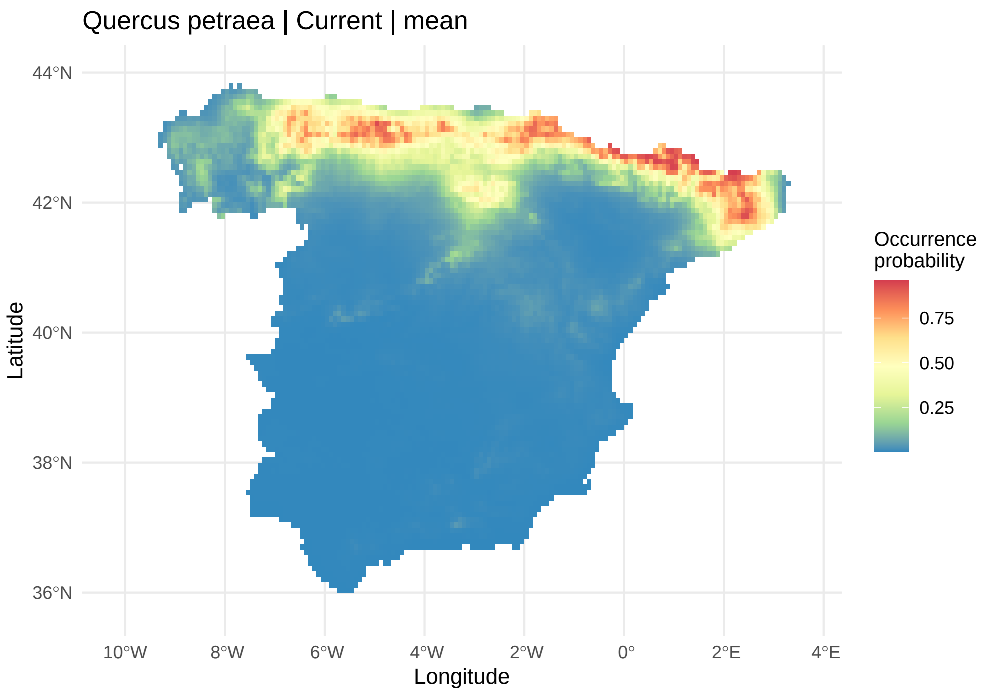

**Fig. S2.2.** Current suitability (posterior mean) for Quercus petraea under the ordered-hierarchical model.

Compare this against Fig. S2.1 above. Constraining the prediction with the global scale typically changes the suitable area relative to the non-spatial baseline.

##### *5.3.2 Prediction uncertainty map*

This shows the posterior standard deviation of the prediction, so you can see where the model is more or less confident. Values are typically higher at range margins or wherever occurrence data are sparse.

```
plot(mod_hierarchical, which = "pred", layer = "sd")
```

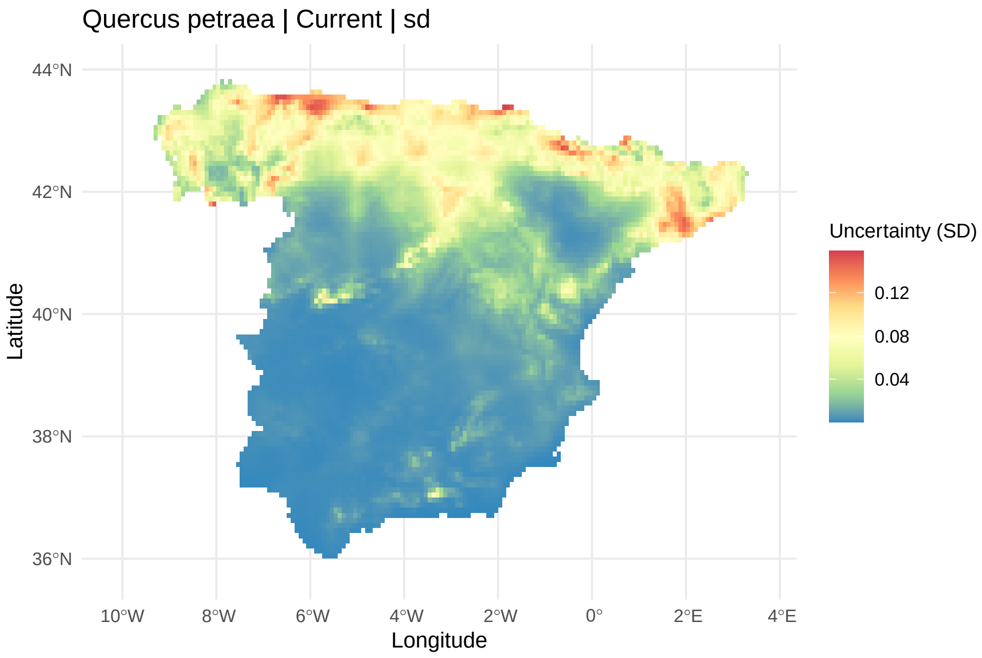

**Fig. S2.3.** Prediction uncertainty (posterior SD). High values indicate areas where the model is less certain, typically at range margins or in regions with sparse occurrence data.

##### 5.3.3 *Future scenario projection*

This shows suitability under the future scenario (`new.env`), using the same fitted model applied to a new set of environmental layers instead of the current ones.

```
plot(mod_hierarchical, which = "Scenario1", layer = "mean")
```

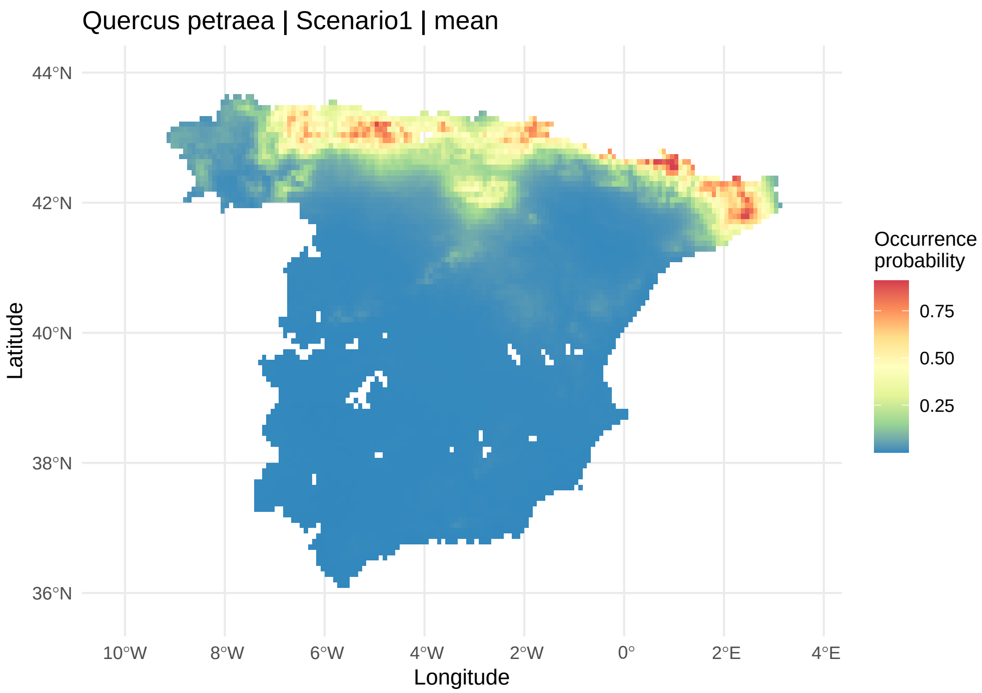

**Fig. S2.4.** Projected suitability (posterior mean) for Quercus petraea under Scenario1.

##### *5.3.4 Spatial random fields*

This shows the two spatial fields the model estimated separately: the broad-scale pattern shared across both datasets ($S_{shared}$), and the fine-scale residual structure specific to the regional data ($S_{RE}$).

```
p_Sshared <- plot(mod_hierarchical, which = "pred_Sshared", layer = "mean")
p_Sre <- plot(mod_hierarchical, which = "pred_Sre", layer = "mean")
patchwork::wrap_plots(p_Sshared, p_Sre, ncol = 1)
```

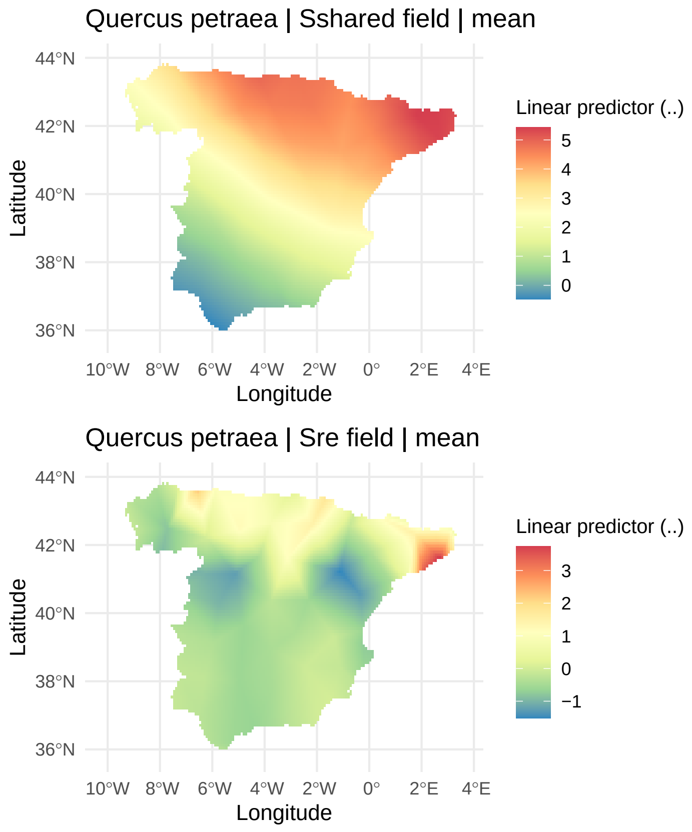

**Fig. S2.5.** Spatial random fields: broad-scale (S shared ) and fine-scale (S RE ).

##### *5.3.5 Residual spatial correlogram*

This shows the residual spatial correlation left in the model's predictions as a function of distance, to check whether the spatial field has captured the autocorrelation it was meant to (a flat line near zero is the goal).

```
plot(mod_hierarchical, which = "correlogram")
```

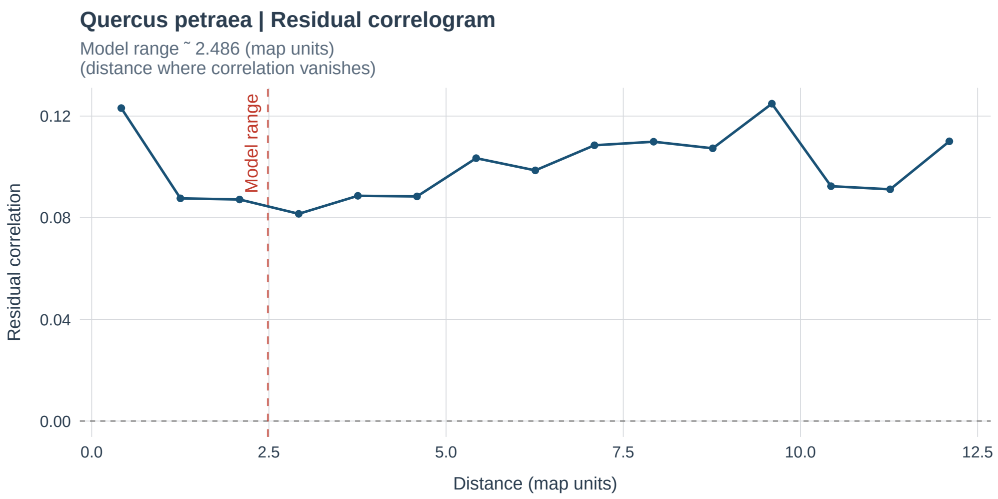

**Fig. S2.6.** Residual spatial correlogram (dashed line = estimated range).

##### *5.3.6 Global vs. regional intercepts*

This compares the posterior distributions of the global and regional intercepts side by side, so you can see how much they overlap, and therefore how much the regional scale is borrowing from the global one.

```
plot(mod_hierarchical, which = "intercepts")
```

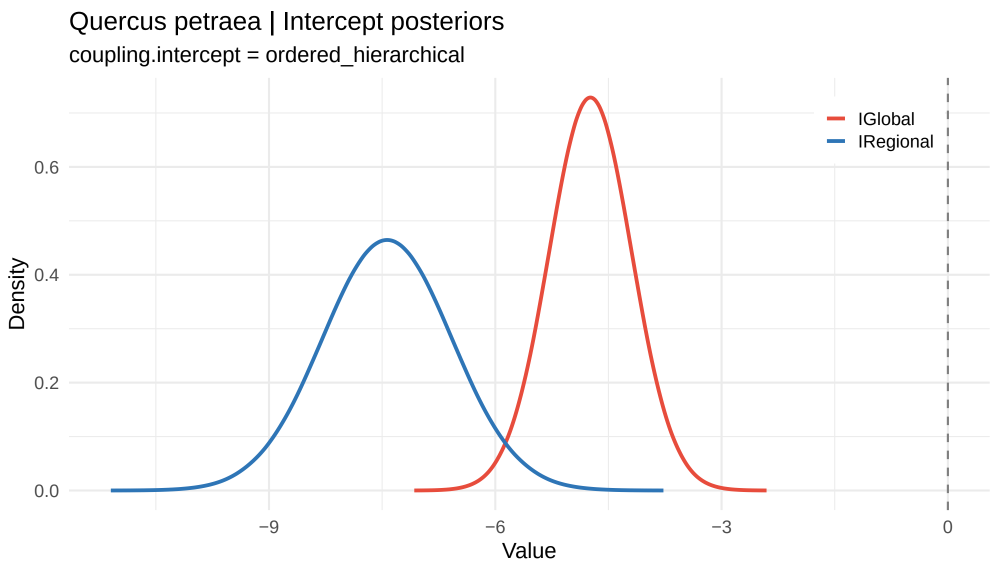

**Fig. S2.7.** Posterior marginal distributions of global and regional intercepts. Overlap indicates borrowing of strength across scales.

### 6 Advanced model configurations

#### 6.1 Non-linear covariate effects

Covariates can be modelled with second-order random walk smoothing (`rw2`) to capture non-linear responses such as unimodal temperature curves. The PC prior `c(u, alpha)` on precision controls the degree of smoothing.

**Note:** non-linear effects pair naturally with `"unpooled"` and `"scale_decomposed"` coupling, since each scale keeps its own independent spline. `"nested_shrinkage"`, `"ordered_hierarchical"`, and `"bayesian_feedback"` instead couple a single linear coefficient across scales, so a non-parametric spline has nothing well-defined to copy or shrink.

```
# u: PC-prior scale, P(spline SD > u) = alpha
# alpha: small alpha = strong belief the response is smooth
covariate_effects <- list(
  regional = list(
    radiation = list(
      model = "rw2",
      u = 0.5,
      alpha = 0.01
    )
  ),
  default = "linear"
)

mod_nonlinear <- MBM.Modelling(
  jmbm_obj                = mySelvars,
  family                  = binomial(link = "logit"),
  spde.mesh               = myMesh,
  regional.pcprior.range = regional.pcprior.range,
  regional.pcprior.sigma = regional.pcprior.sigma,
  shared.pcprior.range    = shared.pcprior.range,
  shared.pcprior.sigma    = shared.pcprior.sigma,
  coupling.intercept      = "unpooled",
  coupling.covariates     = "unpooled", # rw2 requires unpooled predictors
  covariate.effects       = covariate_effects,
  proj.new.env            = FALSE
)

summary(mod_nonlinear)
```

#### 6.2 Cox process formulation

When only presence records are available and background sampling is not desired, the model can be fitted as a log-Gaussian Cox process (LGCP) by setting `family = "cp"`. The output is an intensity surface (expected number of individuals per unit area) rather than an occurrence probability.

**Note:** Any background data contained in `mySelvars` are ignored internally when `family = "cp"`.

```
mod_cp <- MBM.Modelling(
  jmbm_obj                = mySelvars,
  family                  = "cp",          # Log-Gaussian Cox process
  spde.mesh               = myMesh,
  regional.pcprior.range = regional.pcprior.range,
  regional.pcprior.sigma = regional.pcprior.sigma,
  shared.pcprior.range   = shared.pcprior.range,
  shared.pcprior.sigma   = shared.pcprior.sigma,
  coupling.intercept     = "unpooled",
  coupling.covariates    = "unpooled",
  proj.new.env           = TRUE,
  inla.int.strategy      = "eb",
  seed                   = 123
)

summary(mod_cp)
```

---

---

#### 8 Session information

```
sessionInfo()
```

```
## R version 4.3.2 (2023-10-31 ucrt)
## Platform: x86_64-w64-mingw32/x64 (64-bit)
## Running under: Windows 11 x64 (build 26200)
##
## Matrix products: default
##
##
## locale:
## [1] LC_COLLATE=Spanish_Spain.utf8  LC_CTYPE=Spanish_Spain.utf8
## [3] LC_MONETARY=Spanish_Spain.utf8 LC_NUMERIC=C
```

```
## [5] LC_TIME=Spanish_Spain.utf8
##
## time zone: Europe/Paris
## tzcode source: internal
##
## attached base packages:
## [1] stats     graphics  grDevices utils     datasets  methods   base
##
## other attached packages:
## [1] sabinaMBM_0.1.0     INLA_23.09.09       sp_2.2-1
## [4] Matrix_1.6-1.1      inlabru_2.12.0.9015 fmesher_0.3.0
## [7] patchwork_1.3.2     terra_1.8-50        roxygen2_7.3.2
##
## loaded via a namespace (and not attached):
##   [1] DBI_1.3.0               deldir_2.0-4            pROC_1.18.5
##   [4] s2_1.1.7                remotes_2.5.0           testthat_3.2.3
##   [7] rlang_1.1.7             magrittr_2.0.4          otel_0.2.0
##  [10] e1071_1.7-17            tidyterra_0.7.2         compiler_4.3.2
##  [13] callr_3.7.6             vctrs_0.6.5             stringr_1.6.0
##  [16] profvis_0.3.8           pkgconfig_2.0.3         wk_0.9.5
##  [19] fastmap_1.2.0           ellipsis_0.3.2          labeling_0.4.3
##  [22] promises_1.5.0          rmarkdown_2.31          markdown_1.13
##  [25] sessioninfo_1.2.2       ps_1.9.1                purrr_1.0.4
##  [28] xfun_0.52               cachem_1.1.0            jsonlite_2.0.0
##  [31] later_1.4.2             prettyunits_1.2.0       parallel_4.3.2
##  [34] R6_2.6.1                stringi_1.8.7           RColorBrewer_1.1-3
##  [37] parallelly_1.36.0       boot_1.3-28.1           pkgload_1.4.0
##  [40] brio_1.1.5              Rcpp_1.0.14             knitr_1.51
##  [43] future.apply_1.11.2     usethis_2.2.2           httpuv_1.6.16
##  [46] splines_4.3.2           tidyselect_1.2.1        yaml_2.3.10
##  [49] rnaturalearth_1.0.1     rstudioapi_0.17.1       dichromat_2.0-0.1
##  [52] ggtext_0.1.2            codetools_0.2-19        miniUI_0.1.1.1
##  [55] curl_5.2.0              processx_3.8.6          listenv_0.9.1
##  [58] pkgbuild_1.4.7          lattice_0.21-9          tibble_3.2.1
##  [61] plyr_1.8.9              shiny_1.12.1            withr_3.0.2
##  [64] S7_0.2.1                evaluate_1.0.3          future_1.33.1
##  [67] desc_1.4.3              sf_1.0-20               rnaturalearthdata_1.0.0
##  [70] units_0.8-7             spData_2.3.0            proxy_0.4-29
##  [73] urlchecker_1.0.1        xml2_1.5.1              pillar_1.11.1
##  [76] KernSmooth_2.23-22      generics_0.1.4          xopen_1.0.0
##  [79] rprojroot_2.0.4         ggplot2_4.0.3           commonmark_2.0.0
##  [82] scales_1.4.0            globals_0.16.2          xtable_1.8-4
##  [85] class_7.3-22            glue_1.8.0              tools_4.3.2
##  [88] data.table_1.17.2       fs_1.6.6                grid_4.3.2
##  [91] spdep_1.3-3             tidyr_1.3.1             devtools_2.4.5
##  [94] raster_3.6-32           cli_3.6.5               rcmdcheck_1.4.0
##  [97] dplyr_1.1.4             concaveman_1.1.0        gtable_0.3.6
## [100] digest_0.6.37           classInt_0.4-11         htmlwidgets_1.6.4
```

```
## [103] farver_2.1.2           memoise_2.0.1           htmltools_0.5.8.1
## [106] lifecycle_1.0.5        httr_1.4.7              mime_0.13
## [109] gridtext_0.1.5
```
