## Supporting Information S4 for "sabinaMBM: An R package for Multiscale Bayesian species distribution Modelling using INLA"

**Supporting information S4: Extended results from the case study**

**Contents**

S4.1 Robustness of results to background points configuration

Fig. S4.1 Robustness of model performance to background point density

Fig. S4.2 Effect of background sampling strategy on the estimated shared spatial field $S_{shared}$

S4.2 Extended model comparison

S4.3 Extended results: Spatial autocorrelation

Fig. S4.3 Residual spatial autocorrelation for each model configuration

Fig. S4.4 Posterior mean of the random spatial fields estimated by the ordered-hierarchical model

S4.4 Extended results: Scale-dependent covariate effects

Table S4.1 Scale divergence between global and regional covariate effects

S4.5 Future projections and uncertainty

Fig. S4.5 Predictive suitability under current climatic conditions

Fig. S4.6 Multivariate environmental similarity surface (MESS) for future climate projections

Fig. S4.7 Future suitability distributions under hierarchical coupling

Fig. S4.8 Predictive uncertainty under current climatic conditions

Table S4.2 Summary of future habitat suitability predictions across all model configurations

------------------------

### S4.1 Robustness of results to background points configuration

To verify that case study results are not driven by arbitrary choices in background configuration, we evaluated model performance across three background sampling strategies (random, regular grid, spatially stratified) and five-point densities (n = 250, 500, 1,000, 2,500, 5,000; three replicates per density), fitting the non-spatial baseline, spatial regional-only, and ordered-hierarchical configurations for each combination. Across random background densities, the three configurations showed a clear progression of increasing parameter stability with increasing hierarchical complexity (Fig. S4.1). The non-spatial baseline produced consistently poor probabilistic calibration (Brier = 0.109–0.127) and unabsorbed spatial autocorrelation (Moran's I = 0.08–0.13) regardless of background density, confirming that background size does not substitute for spatial structure. Intercept estimates and mean predicted suitability decrease monotonically with background density across all configurations (Fig. S4.1D–E, H); this is an expected structural property of presence-background models, where the intercept absorbs the prevalence ratio and shifts as the number of pseudo-absences increases (Warton & Shepherd, 2010), and does not affect discrimination, calibration, or spatial diagnostics. Both spatial configurations achieved equivalent calibration at n ≥ 1,000 (Brier ≈ 0.065), but the ordered-hierarchical model required n ≥ 1,000 to stabilize its global intercept, spatial field ranges, and scale separation index (SSI > 0.90), with no meaningful improvement at higher densities. Regarding sampling strategy, random background consistently outperformed alternatives in scale separation; regular grid sampling inflated the posterior range of $S_{shared}$ (39.84 vs 18.06 degrees for random), producing an artificially smooth monotonic gradient rather than genuine biogeographic structure (Fig. S4.2), while stratified sampling reduced SSI below 0.80. Random background with n = 1,000 was therefore adopted as the reference configuration for all case study analyses, balancing parameter stability with computational efficiency.

**Fig S4.1** Robustness of model performance to background point density for *Quercus petraea*. Metrics are shown for the ordered-hierarchical (blue), spatial regional-only (orange), and non-spatial baseline (red) configurations across five background densities (n = 250, 500, 1,000, 2,500, 5,000); points and error bars represent the mean ± SE across three replicate background draws. (A) Discrimination capacity (AUC). (B) Probabilistic calibration (Brier score). (C) Posterior field redundancy r² ($S_{RE}$, $S_{shared}$); dashed line marks the 0.95 threshold (ordered-hierarchical only). (D-E) Global and regional intercept ($I_{GL}$ and $I_{RE}$) (log-odds scale; ordered-hierarchical only). (F-G) Posterior range of $S_{shared}$ (ordered-hierarchical only) and $S_{RE}$. (H) Mean relative suitability. (I) Residual spatial autocorrelation (Moran's I).

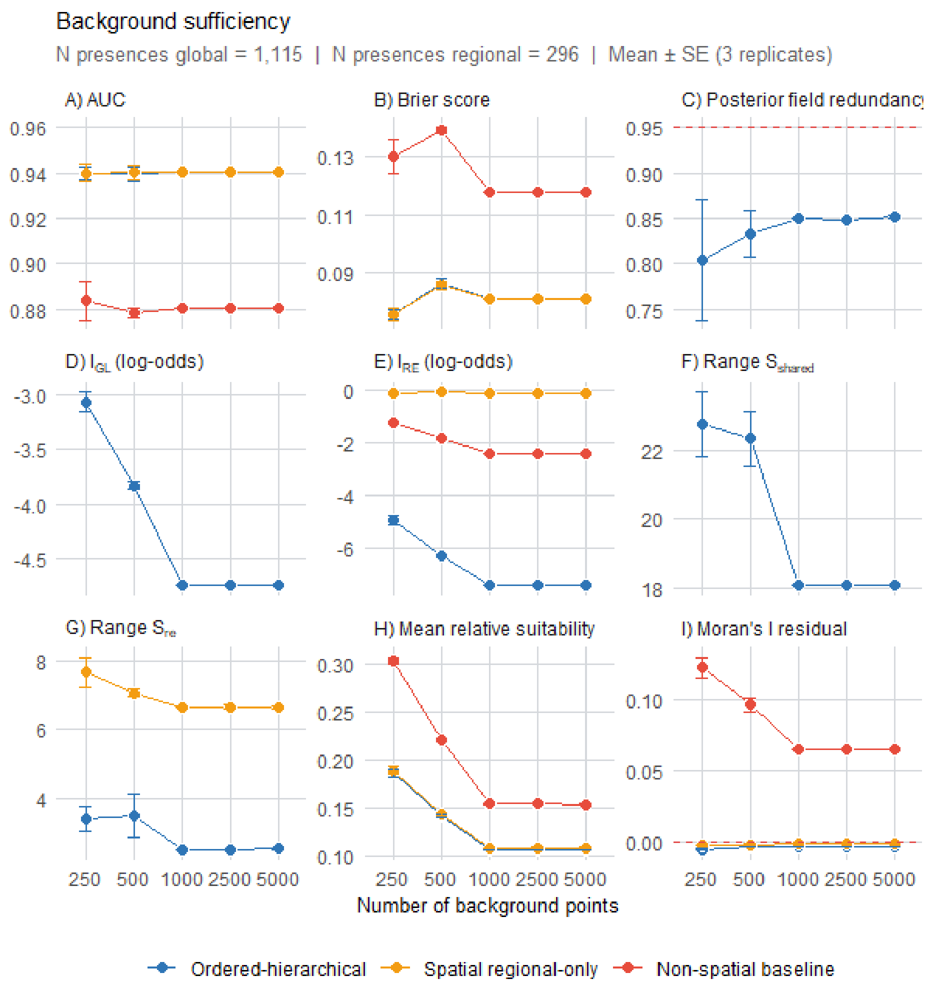


**Fig S4.2** Effect of background sampling strategy on $S_{shared}$ for *Quercus petraea*. (A) Random background (n = 1,000 points; posterior range = 18.1°). (B) Regular grid background (~1,000; posterior range = 39.8°).

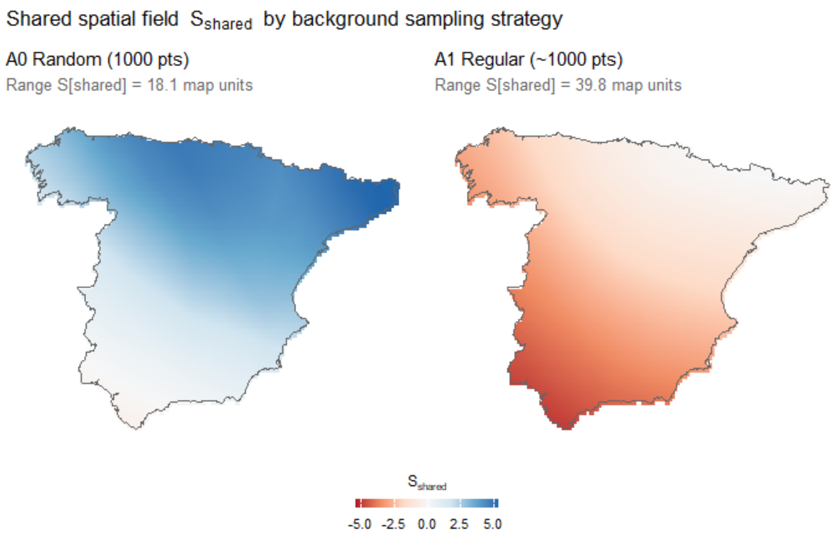

**S4.2 Extended model comparison**

Table 2 (main text) reports full model comparison metrics across all seven configurations. The non-spatial baseline achieved moderate discrimination (AUC = 0.880) with poor probabilistic calibration (Brier = 0.118; log-score = −0.370). Incorporating $S_{RE}$ substantially improved both discrimination (AUC +7%) and calibration (Brier −31%; log-score −28%) over the non-spatial baseline —a valid within-group comparison, since both models use identical regional data. Further integrating global-scale data through multiscale coupling yielded additional log-score improvements over the spatial regional-only model (8–10.5% across joint configurations, peaking with scale-decomposed), with joint multiscale configurations additionally reducing Brier score by up to 5%. Conversely, Bayesian-feedback's log-score gain over the spatial regional-only model remains negligible, despite discrimination and calibration comparable to the other configurations (AUC = 0.939, Brier = 0.082 vs. 0.941 and 0.081 for the spatial regional-only model). In the fully joint configurations, $S_{shared}$ is estimated jointly from both datasets, carrying real cross-scale information into the regional predictions. Bayesian-feedback also transfers $S_{shared}$'s posterior mean and precision per mesh node, rather than re-estimating it from regional data alone.

**S4.3 Extended results: Spatial autocorrelation**

The non-spatial baseline model exhibited strong residual spatial autocorrelation (Moran's I = 0.066), violating the assumption of independent observations. Adding SPDE-based spatial latent fields largely removed this residual autocorrelation across all configurations with active spatial fields (Moran's I = −0.001 to −0.003; Fig. S4.3), improving the reliability of environmental effect estimates. The spatial regional-only model absorbed residual autocorrelation through $S_{RE}$ alone, while the multiscale architectures additionally estimated $S_{shared}$ to capture broad-scale spatial structure simultaneously. To address double-smoothing and weak parameter identifiability inherent in dual-field SPDE models, PC priors enforced structural separation between the range parameters of $S_{shared}$ and $S_{RE}$ (Section 2.4, Supporting Information S1.6). Posterior estimates supported meaningful scale separation: $S_{shared}$ accounted for 57% of spatial variance at a broad scale (posterior range = 18.1 ± 3.0 degrees), while $S_{RE}$ captured the remaining 43% at a fine scale (posterior range = 2.5 ± 1.2 degrees; Fig. S4.4) —an approximately sevenfold difference, consistent with the hypothesis that both broad-scale geographic patterns and finer-scale environmental heterogeneity contribute to the distribution of *Q. petraea* in the

Iberian Peninsula, a decomposition invisible to single-field approaches. Bayesian-feedback shows a different pattern, consistent with its architecture. $S_{shared}$ is transferred from the global fit and held fixed rather than co-estimated with $S_{RE}$.

**Fig. S4.3** Residual spatial autocorrelation across geographic distances, expressed as a Moran's I analogue, for the seven configurations of Table 2. Values near zero indicate successful absorption of spatial structure by the random field/s. The non-spatial baseline serves as reference for the magnitude of unaccounted autocorrelation prior to spatial field inclusion.

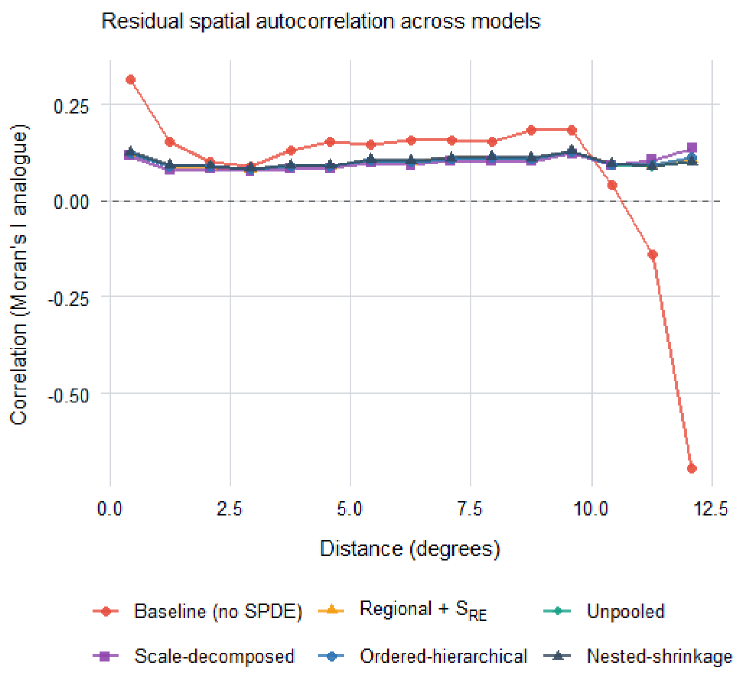


**Fig. S4.4** Posterior mean of the random spatial fields under the ordered-hierarchical model (Table 2). (A) $S_{shared}$ captures broad-scale macroclimatic structure. (B) $S_{RE}$ captures fine-scale spatial heterogeneity. The difference in posterior range reflects the scale separation enforced by the PC priors.

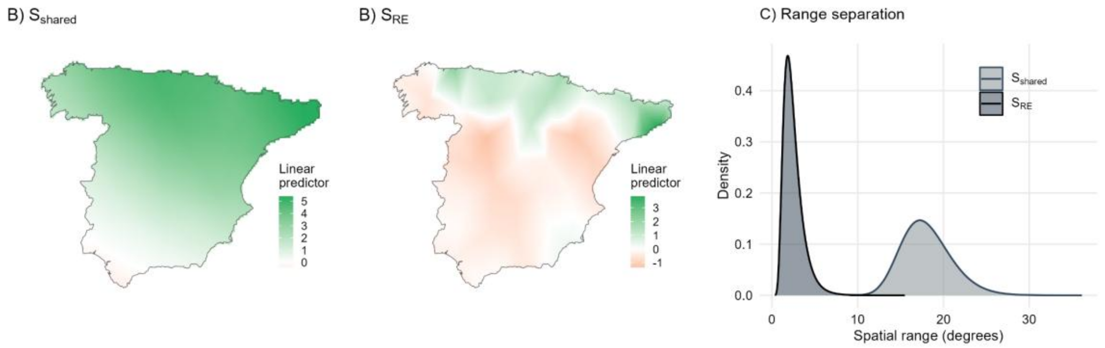


**S4.4 Extended results: Scale-dependent covariate effects**

For annual mean temperature (bio1), ordered-hierarchical shows a consistent negative response at both scales, of greater regional magnitude ($\beta_{GL}$ = -0.086 [95% CI: −0.172, +0.001], $\beta_{RE}$ = -0.241 [−0.378, −0.104]). The global estimate carries substantial uncertainty (its credible interval marginally includes zero) while the regional effect is precisely estimated and credibly negative, suggesting localized intensification of the macroclimatic thermal constraint at the Iberian trailing edge. This pattern is independently retrieved by nested-shrinkage (implied regional effect $\beta_{GL}$ + $\delta_{RE}$ = −0.60 in $\sigma_{RE}$ units; Table S4.1), corroborating that the regional constraint is not an artifact of the ordered-hierarchical parametrization specifically.

In contrast, mean diurnal temperature range (bio2) reveals a scale-specific contrast. At the global scale, the species shows a strong, precise positive association ($\beta_{GL}$ = +0.267 [+0.112, +0.422]), whereas at the Iberian trailing edge, the regional point estimate shifts towards a negative tendency ($\beta_{RE}$ = -0.067 [−0.486, +0.352]). The regional credible interval remains wide, partly reflecting data sparsity at the range margin. This shift, regularized toward the globally-informed niche, suggests that trailing-edge populations may respond uniquely to fine-scale climatic heterogeneity. Notably, this sign reversal is specific to the multiplicative `‘ordered_hierarchical’` mechanism; under `‘nested_shrinkage’`'s additive, zero-centered shrinkage, the regional response remains positive (Table S4.1). This divergence is itself informative. With weak regional signal for bio2, the two hierarchical priors respond differently: `‘ordered_hierarchical’`'s hierarchical copy permits departure from the global sign, while `‘nested_shrinkage’`'s deviation term, shrunk toward zero by a PC-prior, anchors the regional estimate more conservatively to the global trend. This illustrates that coupling architecture choice can materially shape ecological inference for covariates with sparse regional evidence, and should be considered when selecting a configuration for a given system. `‘scale_decomposed’` answers a structurally different question for both variables (global-scale trend vs. regional-scale anomaly rather than global vs. regional total effect) and is not directly comparable coefficient-for-coefficient; full values for all configurations are given in Table S4.1.

**Table S4.1** Scale divergence between global and regional covariate effects for *Quercus petraea*. Positive values indicate a positive association with habitat suitability; negative values indicate avoidance. For the ordered-hierarchical, $\beta_{GL}$ and $\beta_{RE}$ are independently estimated coefficients ($/\sigma_{GL}$ and $/\sigma_{RE}$ respectively). For the scale-decomposed model, $\beta_{GL.res}$ and $\beta_{RE.anom}$ are the broad-scale macroclimatic trend and regional anomaly respectively; both on a common standardization ($/\sigma_{GL}$).

| **Ordered-hierarchical** | | |
|---|---|---|
| **Variable** | $\beta_{GL}$ | $\beta_{RE}$ |
| bio1 | -0.0856 [−0.172, +0.001] | -0.2407 [−0.378, −0.104] |
| bio2 | 0.2673 [+0.112, +0.422] | -0.0670 [−0.486, +0.352] |
| bio12 | 0.0010 [+0.000, +0.002] | 0.0017 [+0.001, +0.003] |
| **Scale-decomposed** | | |
| **Variable** | $\beta_{GL.res}$ | $\beta_{RE.anom}$ |
| bio1 | -0.6532 [−0.877, −0.430] | -0.0914 [−0.305, +0.123] |
| bio2 | 0.1016 [−0.489, +0.692] | 0.1817 [−0.705, +1.068] |
| bio12 | 0.0030 [+0.001, +0.005] | -0.0008 [−0.003, +0.001] |

#### S4.5 Future projections and uncertainty

The global prior embedded in the ordered-hierarchical configuration is designed not to distort predictions where regional data are informative —confirmed by the equivalent current performance across configurations (Table 2; Fig. S4.5)— but its value emerges under future projection, where the regional model is asked to predict beyond its calibration domain. Under the 2070 SSP585 scenario, the structural consequences of niche truncation become evident. Despite future climates remaining broadly analogous to the current calibration space across most of the Spanish domain (Fig. S4.6), the regional-only model predicts greater suitable habitat extent than its hierarchical counterpart. A larger proportion of the Peninsular area is identified as potentially suitable above the decision boundary where the linear predictor crosses zero (equivalent to p = 0.50 on the response scale; 5.7% vs 4.3%; Fig. 3A-B cf Fig. S4.7; Table S4.2). This divergence does not arise from projecting into climatically novel conditions, but from a fundamental limitation of calibrating at range margins (Scherrer et al., 2021). The Iberian calibration space captures only 74% of the global niche, meaning that the regional model cannot estimate responses to the climatic conditions the species regularly avoids across its full distribution. Without exposure to those conditions during calibration, the regional model is structurally unable to self-constrain its future predictions. Whether this leads to overprediction remains unresolved without further empirical validation, but the hierarchical model's superior out-of-sample discrimination in data-poor regions indicates that the additional constraint is ecologically informative rather than artificially conservative.

The posterior distribution of $\beta_{copy}$ (mean = 1.578, 95% CI [1.225, 1.940], well above and clearly separated from the prior centred at 1; Fig. 3D), indicates that regional data strongly amplify the global baseline. This demonstrates that global-scale information contributes substantially to regional inference, as expected for range-margin populations where regional data alone fail to capture broader environmental relationships. Spatially, this hierarchical constraint systematically reduces future suitability across the northern range, the area of highest species occurrence and greatest model divergence (Fig. 3C). Consequently, mean future suitability decreases by ~16% (from 0.089 to 0.0759; Table S4.2). Spatial cross-validation (training models on two regions and evaluating on held-out records from the third) confirms this advantage is localized where regional data are scarcest. In the data-poor trailing-edge region (Region 3, n = 26), the ordered-hierarchical model substantially improves out-of-sample discrimination (AUC = 0.963 vs 0.693 for the regional-only), and reduces the Brier score by 71.3%. Conversely, in data-rich northern regions (e.g., Region 2, n = 119), performance differences are negligible (ΔAUC = +0.009; Fig. 4). This confirms that hierarchical constraint may act as a safeguard where regional evidence is insufficient to self-constrain, and that its value is empirically verifiable through held-out validation rather than merely theoretical.

Figure S4.7 shows the full distributional consequence of this constraint across all Iberian pixels, on a logit-transformed scale, a representation that compresses the extremes near 0% and 100% and expands the intermediate range where habitat suitability assessments are most consequential. The regional-only distribution presents a long right tail extending well beyond the 50% threshold, consistent with predictions generated under niche truncation, where the regional model lacks exposure to the climatic conditions the species avoids globally. The ordered-hierarchical distribution preserves the same general shape, confirming that the prior does not distort predictions across the bulk of the distribution where regional data are informative, but with the right tail substantially compressed. Simultaneously, the hierarchical framework concentrates significantly more probability mass near zero. This demonstrates that incorporating global distribution limits prevents over-extrapolation in highly suitable areas while increasing model certainty when discounting unsuitable habitats. The 50% threshold, beyond which a pixel is considered potentially suitable habitat, makes this compression directly interpretable for conservation planning, as the hierarchical framework substantially reduces the extent of predicted suitable habitat by compressing the high-suitability tail that regional calibration alone cannot constrain.

Beyond constraining point predictions, the hierarchical model reduces mean posterior standard deviation by 24.3% relative to the regional-only counterpart (mean SD = 0.026 vs 0.034), with lower uncertainty in 77.4% of pixels, yet this global average conceals a spatially structured and ecologically interpretable pattern (Fig. 5 cf Fig. S4.8). In the northern fringe, where *Q. petraea* maintains all its Iberian populations, the hierarchical model simultaneously reduces future suitability estimates and posterior uncertainty relative to the regional-only counterpart (Fig. 3C and 5C). The positive spatial correlation between ΔSuitability and ΔSD (r = +0.18) reveals that the constraint acts coherently across both quantities in the same locations. Where the global prior most strongly revises suitability downward, it also most strongly reduces predictive uncertainty. This is corroborated by the reduction in mean credible interval width (0.100 vs 0.131 for the regional-only model), and by the spatial congruence between Figs. 2C and 4C, where the areas of greatest suitability revision and greatest uncertainty reduction overlap geographically along the northern margin. A regional-only model would have reported high suitability with artificially wide posterior intervals in these areas, conveying unwarranted imprecision about predictions that global information constrains.

**Fig. S4.5** Predictive suitability under current climatic conditions for *Quercus petraea*. (A) Spatial regional-only and (B) ordered-hierarchical model (Table 2). (C) Pixel-wise difference (ΔSuitability = regional - hierarchical); warm colors indicate more conservative predictions; values near zero indicate agreement.

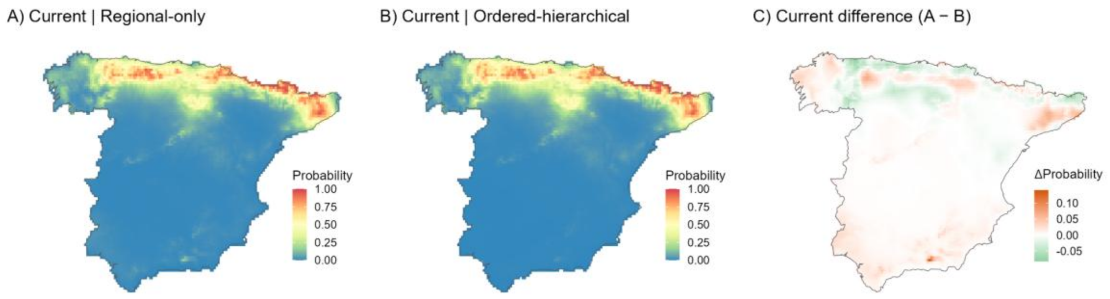


**Fig. S4.6** Multivariate environmental similarity surface (MESS) for future climate projections, computed relative to the regional calibration space. Positive values indicate future climates analogous to conditions within the calibration envelope; negative values indicate partial extrapolation beyond the range of calibration conditions.

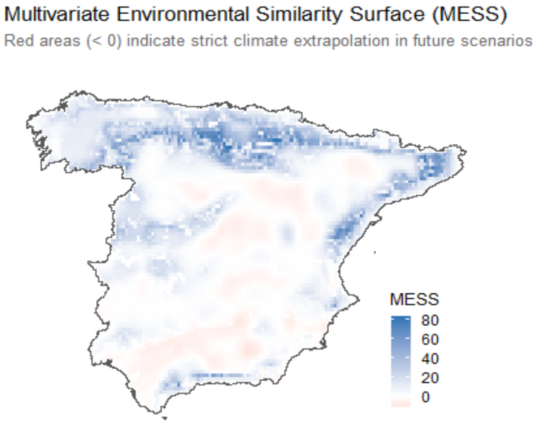


**Fig. S4.7** Future suitability distributions under hierarchical coupling for *Quercus petraea*. Kernel density of predicted suitability for the regional-only (red) and ordered-hierarchical (blue) models (Table 2), on a logit-transformed scale and suitability-equivalent axis labels. The dashed line marks the logit = 0 decision boundary (p = 0.50), beyond which a pixel is considered potentially suitable.

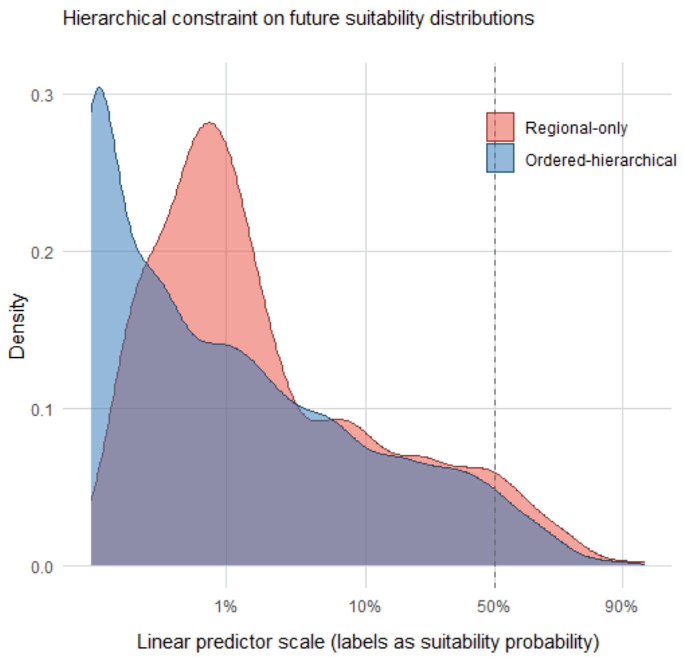


**Fig. S4.8** Predictive uncertainty under current climatic conditions for *Quercus petraea*. Posterior standard deviation (SD) of the (A) regional-only and (B) ordered-hierarchical model (Table 2). (C) Pixel-wise difference in predictive uncertainty (ΔSD = $SD_{regional}$ - $SD_{hierarchical}$); warm colours indicate lower uncertainty under the hierarchical model; cool colors marginally greater uncertainty.

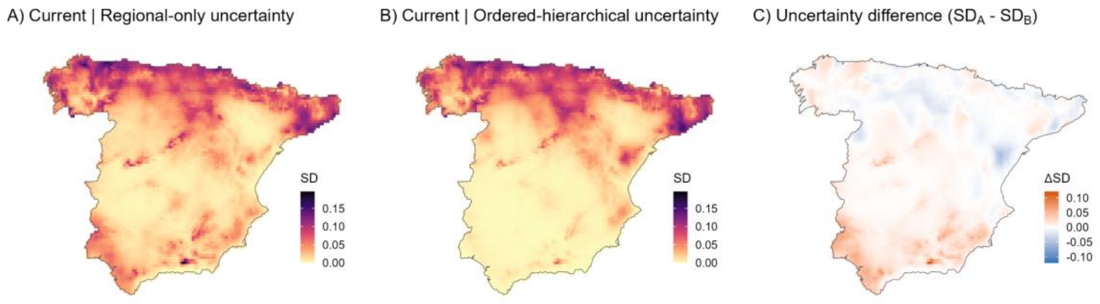


**Table S4.2** Summary of future habitat suitability predictions across all model configurations for *Quercus petraea*. Mean suitability, 90th percentile (P90), and fraction of the Iberian Peninsula predicted above the 50% suitability threshold are reported for each configuration.

| Model | Mean suitability | P90 | Fraction >50% |
|---|---|---|---|

| | | | |
|---|---|---|---|
| Non-spatial baseline | 0.070 | 0.213 | 1.6% |
| Spatial regional-only | 0.089 | 0.340 | 5.7% |
| Unpooled | 0.087 | 0.329 | 5.4% |
| Ordered-hierarchical | 0.075 | 0.285 | 4.3% |
| Nested-shrinkage | 0.082 | 0.319 | 5.1% |
| Scale-decomposed | 0.051 | 0.186 | 1.6% |
| Bayesian-feedback | 0.094 | 0.362 | 6.2% |
